# In situ structures of the ciliary tip show the multi-step mechanism of axoneme assembly

**DOI:** 10.64898/2026.01.30.702858

**Authors:** Helen E. Foster, Samuel E. Lacey, H. Noor Verwei, Margot Riggi, Jose Davila-Velderrain, Gaia Pigino

## Abstract

Motile cilia are highly complex, ordered organelles which extend from the cell surface to generate fluid flow. During motile cilia assembly, components rapidly self-organize at the ciliary tip. Here, over two hundred different components bind microtubules to form an intricate, repeating structure called the axoneme. Despite in-depth structural characterization of axonemal complexes, we have limited understanding of how these structures are generated. Here, we directly visualize the steps of axonemal assembly *in situ* by performing cryo-electron tomography of *Chlamydomonas reinhardtii* cilia tips. Using subtomogram averaging and classification, we identified conserved microtubule luminal proteins that establish periodicity and show that controlled microtubule lattice perforation can facilitate incorporation of late-binding components. By uncovering unexpected assembly intermediates, we explain how structural integrity and the emergence of motility are coordinated within the axoneme. Overall, we demonstrate that self-assembly proceeds through regulated, sequential construction steps and provide a molecular framework for understanding axoneme biogenesis.

## Introduction

Motile cilia (also called flagella in some species) are highly complex organelles that protrude from the surface of specialized eukaryotic cells and beat to drive fluid flow or provide cellular propulsion. Powerful, repetitive ciliary beating relies on careful positioning of specialized dynein motors and regulatory complexes within the axoneme, the highly ordered microtubule bundle found at the core of motile cilia. Recent advances in cryo-electron microscopy have shown the intricate placement of over 200 distinct, periodically repeating components on axonemal microtubules ^1–4^, and how these differ in different organisms ^1,5–8^ or cell types ^9^. Defects in axoneme formation leads to a broad-spectrum of congenital conditions called ciliopathies ^10^, however the process of high fidelity ciliary assembly is poorly understood.

Motile axonemes consist of nine doublet microtubules (DMTs) that each consist of a 10-protofilament B-tubule tethered to the side of a 13-protofilament A-tubule, surrounding a central pair (CP) of singlet microtubules. Axonemes are initially templated by the basal body, with DMTs directly extending from the ends of basal body triplet microtubules ^11,12^ and the CP being independently nucleated ^12–14^. Axoneme elongation proceeds through polymerization of microtubule plus ends, which are clustered close to the distal end of the cilium, in a region called the tip ^15–17^. The tip constitutes a distinct ciliary sub-compartment and is characterized by a unique axonemal morphology and protein composition compared to the shaft ^12,13,18–20^. Highlighting this, DMT growth is staggered at the tip, with B-tubules terminating up to 1 micron from the cilium distal end and A-tubules extending alone as singlets.

Both DMTs and the CPs are coated with periodically repeating protein complexes that create and modulate the ciliary beat. It is thought that a sequential, hierarchical binding of MT-associated proteins templates these repeating structures during assembly. However, identifying the foundational factors that establish this periodicity has proven difficult through analysis of fully assembled structures. While differences between nascent and mature doublets exist ^21^, we lack a molecular explanation for axoneme biogenesis, and the roles of each individual protein to ciliary assembly. Moreover, the proteins associated with CP microtubules to form the central apparatus (CA), are nearly completely unique compared to those on DMTs, suggesting a distinct assembly process takes place. In addition, it is unclear whether axoneme assembly is achieved autonomously, solely using components within the axonemal repeats, or whether other tip-specific assembly factors are involved.

To understand how axonemes are generated, we performed cryo-electron tomography (cryo-ET) of Chlamydomonas reinhardtii cilia tips, capturing both DMTs and the CA in distinct stages of assembly. Through subtomogram averaging and classification, we found that DMT assembly comprises five stages. We identified the highly conserved microtubule inner proteins (MIPs) FAP53 and FAP127 as pioneers which template the DMT repeat, and discovered that non-canonical assembly intermediates form during initial B-tubule formation. Unexpectedly, we found that temporally and spatially controlled perforation of the microtubule lattice occurs during assembly, which may facilitate late incorporation of luminal microtubule inner proteins (MIPs). Furthermore, we show that the assembly of DMTs and CA is not confined to the tip region, but it extends proximally since several axonemal components are only present in the mature axoneme. Our findings demonstrate that axoneme assembly involves a tightly regulated, modular process which requires tip-specific factors.

## Results

### Steady-state cilia are in state of suspended growth

To investigate changes in axonemal structure during different phases of assembly, we imaged Chlamydomonas ciliary tips by cryo-ET in three conditions (Figure 1A-B); steady state (SS), regrowth (Rg), and resorption (Rs). We compared the tip architecture across all conditions by modelling the microtubule morphology and mapping their visible decorations, defining three DMT sections for analysis, namely singlet, doublet noODAs, doublet withODAs (Figure 1C, E1A). Singlet and doublet noODAs lengths were similar in regrowing and steady-state conditions, but shortened during cilia resorption (Figure 1D, E1B). Interestingly, we found ODAs bound to singlet A-tubules only during resorption (Figure E1B), indicating this is not merely the reverse of assembly and involves distinct, depolymerization-specific steps.

**Figure 1.**
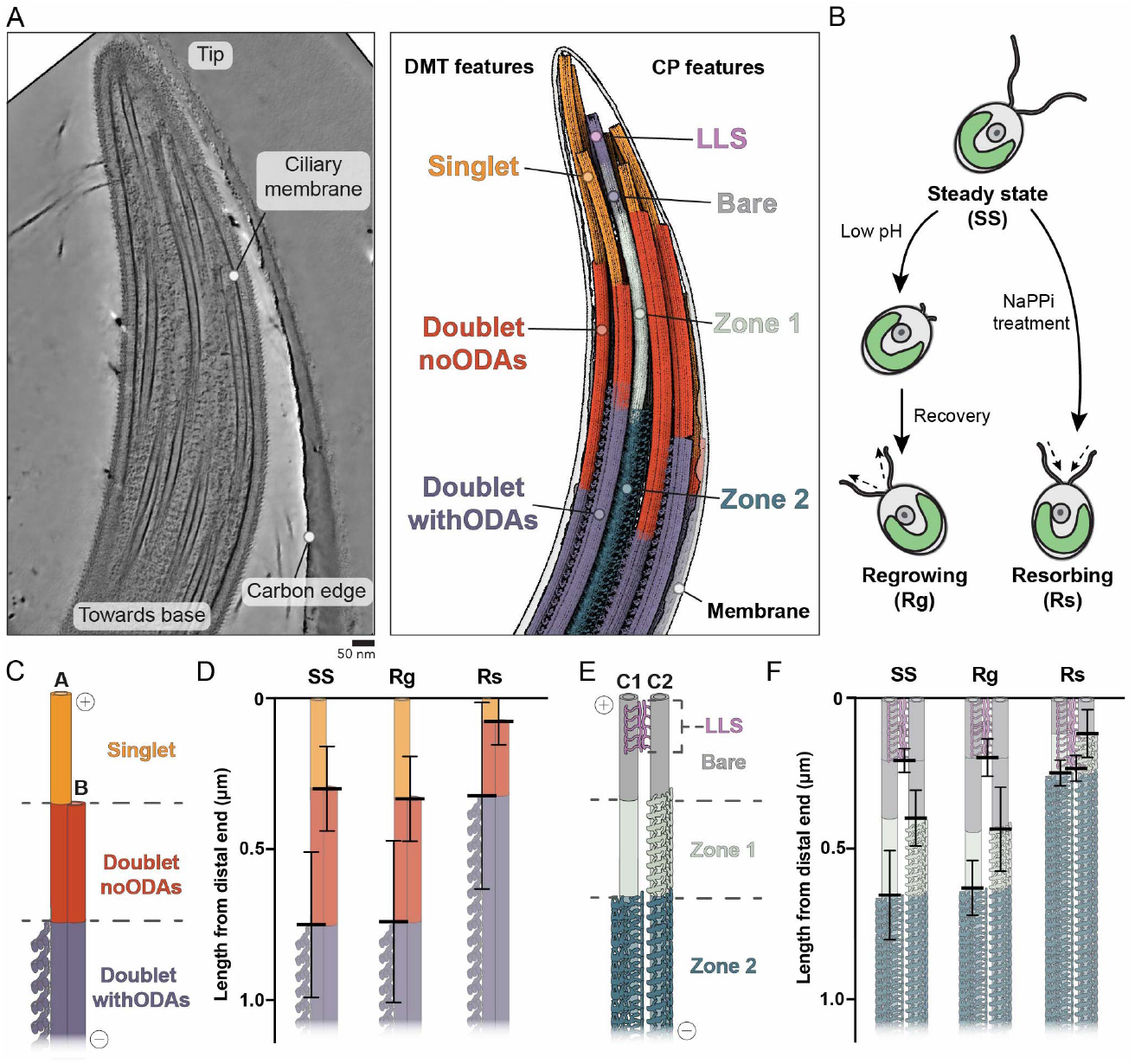
Steady state and regrowing ciliary tips show similar morphology. A) Changes in microtubule architecture at the Chlamydomonas reinhardtii ciliary tip. Left: example tomogram of SS cilia tip. Right: Corresponding 3D model showing annotated features of doublet microtubules (DMTs) and central pair (CP). B) Schematic illustrating how motile cilia were induced to transition from steady state (SS) to either Regrowing (Rg) or Resorbing (Rs) conditions. C) Model of the three DMT sections observed at the cilia tip. From the microtubule plus ends, a singlet A-tubule gains the B-tubule to form a doublet then the outer dynein arms (ODAs) are added. D) Stacked bar chart showing DMT section lengths per microtubule from distal (plus) end. Mean ± SD shown. Statistical comparisons are provided in Figure E1B. E) As in (C), but for the CP. The LLS is bound at the plus end, followed by three additional zones of microtubule decoration. F) As in (D) but for the CP, with statistical analysis in Figure E1D.

The CA showed four zones with distinct morphology in the tip (Figure 1E). We found the previously described ladder-like structure (LLS) positioned between the two CP MTs at their plus-ends ^12,14,22^ (Figure E1C). The LLS maintained similar length in all conditions (Figure 1F, E1D), tracking the microtubule plus-ends during growth and depolymerization. Proximal to the LLS, there was a *bare* region with no visible repeating features on either MT. Next, only one MT had repeating structures on its surface (*Zone 1*) and subsequently both MTs gained decoration typical of the CA ^1,2,23^ (*Zone 2*; Figure E1E). The length of CP sections was similar in SS and Rg tips but significantly shorter in Rs conditions (Figure 1F, E1D). Overall, both DMT and CP showed similar morphology during steady state and regrowth, indicating comparable processes occur in these conditions, possibly with different kinetics. Conversely, we find that resorption is characterized by a distinct process.

Steady state cilia are proposed to exist in a state of dynamic equilibrium ^24,25^ which allows constant fine-tuning of mature cilia length ^26–30^. The similar morphology of microtubules within SS and Rg tips supports this hypothesis and suggests the SS condition represents a suspended state of growth. We therefore analyzed axonemal structures within both SS and Rg cilia tips to uncover the molecular mechanism of axoneme assembly.

### Mapping out the DMT assembly process

An undecorated, nascent singlet microtubule has an intrinsic 8-nm repeat dictated by the tubulin lattice. In contrast, fully assembled DMTs are coated with over 120 different proteins which are organized on the internal and external surfaces of A- and B-tubules with higher order periodicities of 8, 16, 24, 48 or 96 nm ^3,4^. We set out to understand how these higher-order periodicities, B-tubule formation, and the binding of motility machinery are established.

To resolve DMT assembly intermediates, we generated subtomogram averages of each microtubule segment (*singlet, doublet noODAs* and *doublet withODAs*) and compared these to averages of the axoneme shaft, which is the *mature* structure. Iterative refinement and classification (detailed in Supplementary Information) revealed progressive incorporation of known axonemal components and uncovered novel, stage-specific structural features. Based on these data, we define five sequential assembly stages (Stages 1-5) that provide a framework for DMT assembly.

### Stage 1: The filamentous MIPs FAP53 and FAP127 are pioneer factors

The singlet region at the tip represents the earliest stage of DMT assembly. To determine whether higher-order periodicities beyond the 8 nm tubulin repeat are already established at this stage, we analyzed Stage 1 subtomogram averages. These lacked large axonemal complexes, but showed two small densities on the inner surface of the SS *singlet* (Figure 2A-B, E2A). Based on their morphology and position between protofilaments A06-A07 and A07-A08, we identified these densities as the filamentous MIPs (fMIPs) FAP53 (mammalian CCDC11) and FAP127 (mammalian MNS1). In the Rg *singlet*, these densities were smeared and not well resolved (Figure E2B), indicating partial occupancy and suggesting that FAP53 and FAP127 are added soon after microtubule formation rather than co-polymerizing. Thus, FAP53 and FAP127 are the earliest proteins to bind nascent microtubules at the ciliary tip, acting as the pioneering factors for DMT assembly. Both FAP53 and FAP127 are known to assemble end-on-end with 48 nm periodicity ^3^ and are therefore capable of templating higher order axonemal repeats. Notably, both proteins extend an α-helix through the A-tubule lattice between protofilaments A10-A11, corresponding to the outer junction (OJ) ^31^, the site of B-tubule nucleation. FAP53 and FAP127 can thus pattern the A-tubule longitudinal repeat and define the future site of B-tubule formation. This shows that the internal 48-nm patterning precedes the establishment of external 96-nm repeats during DMT assembly.

**Figure 2.**
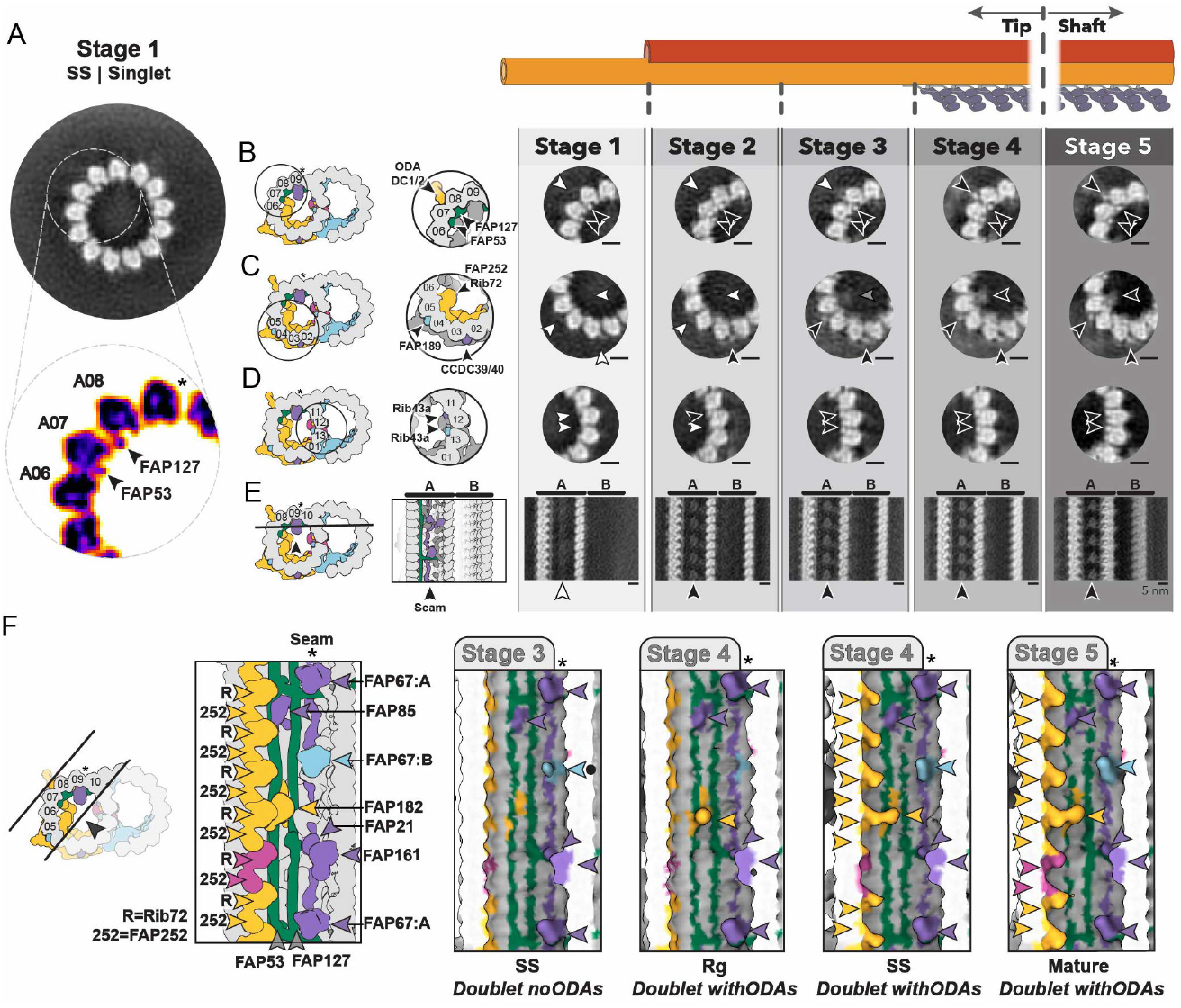
Luminal fMIPs and seam binding factors initiate DMT assembly. A) Singlet microtubules contains the fMIPs FAP53 (between A06-A07) and FAP127 (between A07-A08). Projection of subtomogram average (8 nm repeat, SS tips) in greyscale (top panel) and ImageJ ‘Fire’ LUT (inset below). B-E) Subtomogram average projections from each stage of DMT assembly in SS cilia tips and the *mature* axoneme. Left: schematic of selected doublet region. Right: cartoon highlighting components expected based on PDB 8GLV. Arrowheads indicate assignment of protein occupancy - absent (white), partial occupancy (grey) and or present (black). B) fMIPs FAP53 and fMIP127 are present in all stages, ODA-DC appears in Stage 4. C) FAP59/172 (mammalian CCDC39/40) appears from Stage 2, FAP189 from Stage 3 and luminal MIPs FAP252/Rib72 from Stage 4. D) Rib43a binds between A11-A12 in Stage 2 and between A012-13 in Stage 3. E) Seam-binding proteins are present from Stage 2 onward. F) Surface view of 96 nm repeat showing progressive addition of seam-binding proteins and luminal MIPs. Proteins consistently present are (FAP161, FAP21 and one copy of FAP67 (FAP67:A); purple) along with FAP85. A second copy of FAP67 (FAP67:B, light blue) is weakly present in SS Stage 3 (black dot) and absent from Rg Stage 4. FAP182 binds in Stage 4. Asterisks indicate seam position.

### Stage 2: Addition of seam-binding proteins triggers B-tubule assembly

To identify factors that initiate B-tubule formation, we compared Stage 1 and Stage 2 averages and found three densities that are added concurrently with the singlet-to-doublet transition. These map to: 1) the outer A-tubule surface between A02-A03 (Figure 2C, E2C), occupied by FAP59/172 (mammalian CCDC39/40), a key determinant of 96-nm component positioning ^4,32^, 2) the A-tubule inner wall at A11-A12 (Figure 2D, E2D), occupied by Rib43a and 3) the inner A-tubule surface close to the seam (Figure 2E, E2E), a structural landmark for microtubule orientation ^33,34^.

Additional analysis showed that among all the seam-binding proteins, FAP161, FAP21, FAP85 and only one copy of FAP67 (mammalian NME7) were consistently detected across all steady-state and regrowing averages (Figure 2F), whereas the second copy of FAP67 was absent until Stage 5. This indicates that DMT assembly proceeds through sequential generation of binding sites rather than simultaneous recruitment of components available at the tip.

To test whether FAP59/172, Rib43a or the seam-binding components promote B-tubule formation by remodeling the microtubule lattice architecture, we analyzed inter-protofilament angles. In vitro, singlet microtubules are roughly cylindrical ^35^ but doublet microtubules exhibit pronounced lattice deformation ^7,36,37^. We found the transition from Stage 1 to Stage 2 of assembly is marked by a series of structural changes in the lattice (Figure E2F). The major change was a local increase in curvature at the seam, while the Rib43a and FAP59/172 binding sites remaining unchanged. These results indicate that although not all seam-binding proteins are required, FAP67, FAP21, FAP161 and FAP85 are key for locally remodeling the A-tubule lattice to initiate B-tubule formation.

### Stages 2-3: Inner junction closure does not rely on the presence of PACRG

We next asked how B-tubule closure is achieved at the inner junction (IJ). In a mature axoneme, FAP20 and PACRG form the 8-nm repeating ‘IJ filament’ ^3,31^ which bridges the A- and B-tubules between A01 and B09^3,31,38,39^. The IJ is further fortified by FAP52, FAP276 and FAP106 (ENKUR in mammals), which link the B-tubule inner wall to the A-tubule outer wall in a 16-nm repeating structure, and by the fMIPs FAP210 and FAP45, which line the B-tubule lumen near the IJ^38,40,41^. Proper IJ formation is essential for mechanical stability of DMTs ^38,40–43^ but the specific role of these individual components in IJ assembly and their assembly logic remained unclear.

The analysis of our data shows that IJ formation proceeds via a unique, tip-specific non-canonical intermediate. During initial B-tubule formation (in Stage 2), the IJ contains an uncharacterized structure with 8 nm repeat (Figure 3A, E3A) that does not resemble canonical IJ components, suggesting the presence of a transient scaffold responsible for initial IJ closure. In Stage 3, this scaffold is replaced by the canonical 16-nm repeating FAP52/106/276 structure (Figure 3A, E3A-B), coincident with binding of the fMIPs FAP45 and FAP210 (Figure 3B, E3C). Our analysis of inter-protofilament angles showed that the A-tubule lattice changes concurrently with inner IJ conversion in Stage 3: the wall between A- and B-tubules flattens and the A-tubule near the IJ gains high curvature (Figure E2F). These features approach the final conformation, indicating that DMT maturation is incomplete prior to canonical IJ formation.

**Figure 3.**
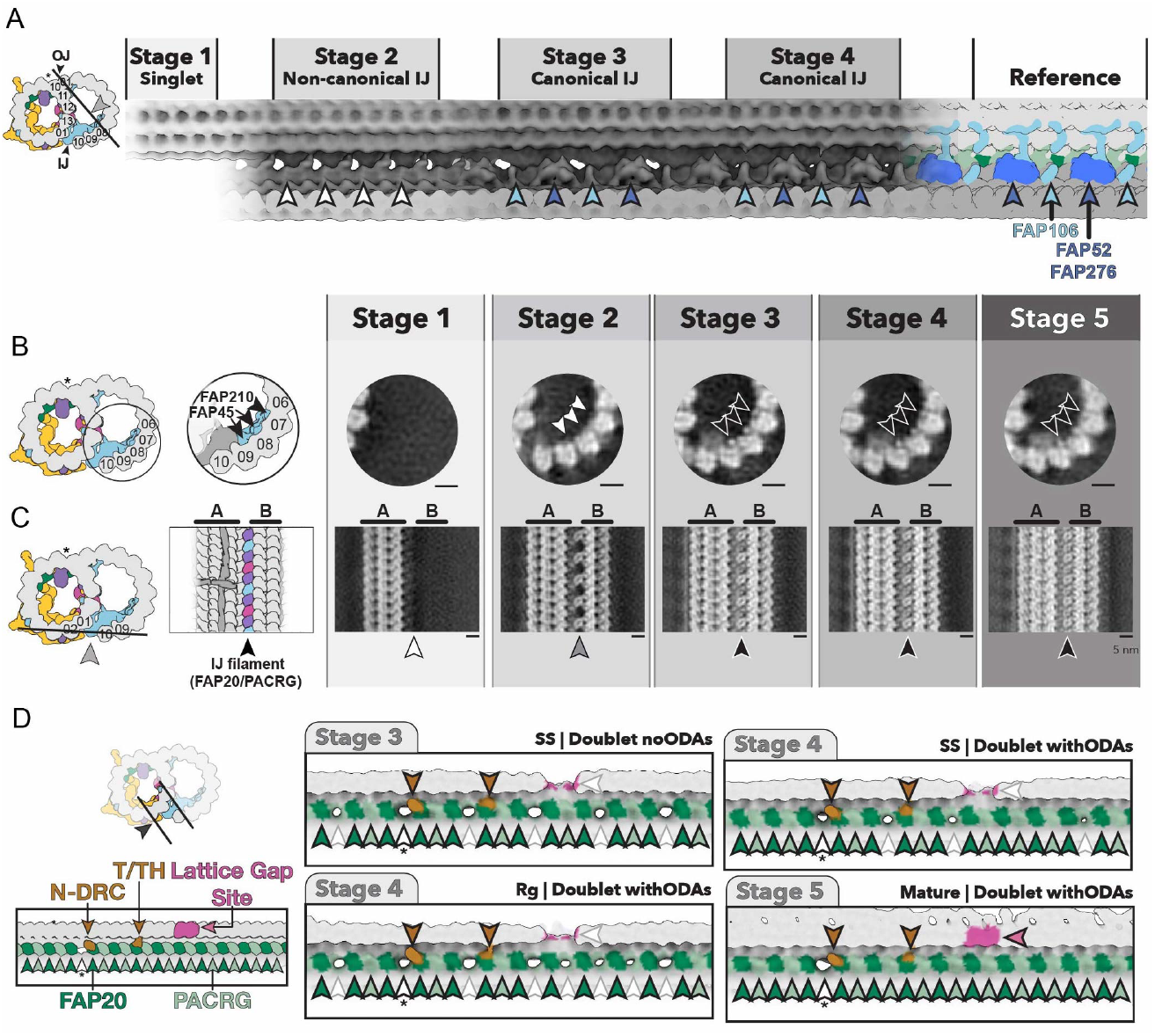
The IJ matures through a non-canonical intermediate. A) Structural transitions at the internal face of the IJ in SS cilia tips. Stages 1-2: 8 nm repeat averages; Stages 3-4: 16 nm repeat averages. Far-left: cartoon showing viewing perspective. Far-right: expected appearance of components based on PDB 8GLV. Arrowheads indicate presence of FAP52/FAP276 (dark blue), FAP106 (light blue) or component absence (white). B-C) Subtomogram average projections from each stage of DMT assembly in SS cilia tips and the *mature* axoneme. Cartoons indicate viewing perspective and expected appearance of components. Arrowheads indicate assignment of protein occupancy - absent (white), partial occupancy (grey) and or present (black). B) B-tubule fMIPs FAP210 and FAP45 bind at Stage 3, concurrently with IJ conversion from a non-canonical to canonical state. C) The IJ filament progressively forms throughout Stages 2-4. D) Variable PACRG occupancy is seen throughout assembly in the 96 nm repeat averages. Absent components indicated with white arrowheads, asterisk indicates site known to lack PACRG in mature axoneme.

Surprisingly, the IJ filament itself (containing FAP20/PACRG) remains incomplete throughout DMT assembly. In Stage 2, FAP20 was present, but PACRG absent (Figure 3C, E3D). Across different assembly stages in SS and Rg tips, PACRG showed only partial occupancy (Figure 3D) and this was not strictly linked to FAP52/106/276 binding (Figure E3E). PACRG incorporation is therefore not required for B-tubule closure and occurs separately from other IJ components. Variable occupancy of PACRG was previously described in various organisms (Figure E3F). In Trypanosoma, this reflects two paralogous specialization (PACRG-A and PACRG-B^44,45^). However, *Chlamydomonas* express a single PACRG^46^, yet shows preferential occupancy at the ‘PACRG-A site’ over the ‘PACRG-B site’ during assembly, indicating differential occupancy must arise via another mechanism.

### Stages 3-4: Microtubule lattice gaps facilitate assembly

Many MIPs are absent in Stages 1-3 and are incorporated only at Stage 4, including Rib72, FAP252 and FAP115, which bind the A-tubule inner wall (Figure 2C,F). Their late addition raises the question of how they access the microtubule lumen. Although they could theoretically diffuse ∼700 nm from the microtubule plus-ends (Figure 1D), the A-tubule ends are obstructed by plug-like structures throughout the cilia regeneration ^14,47^ (Figure 4A, E4A-B).

**Figure 4.**
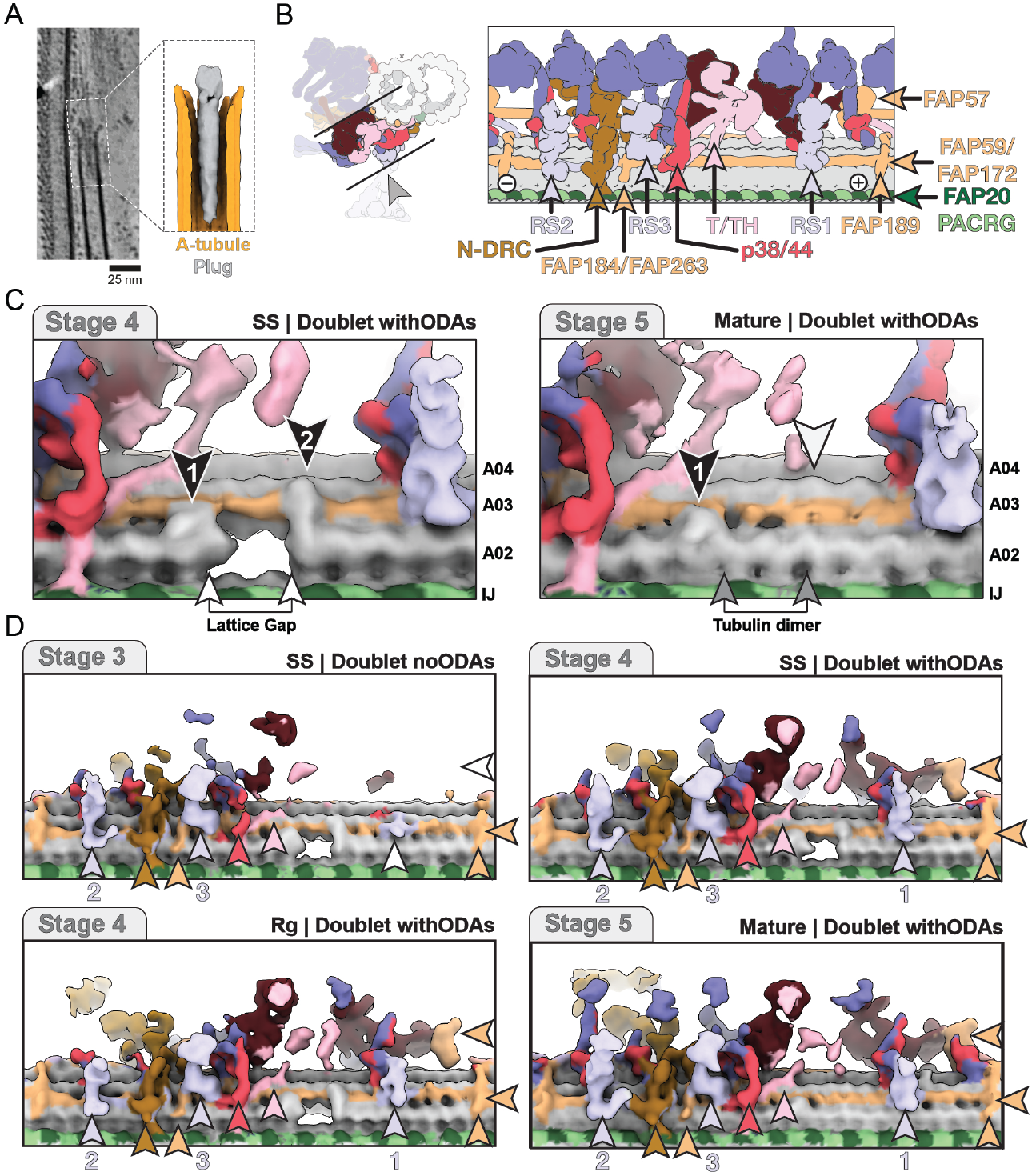
Tip-specific lattice gaps can facilitate assembly. A) Slice through tomogram showing example A-tubule plug. Subtomogram average of A-tubule plugs from SS cilia is shown in inset. B) Cartoons showing viewing perspective and location of key components in 96 nm repeat indicated in C-D C) Position of tubulin-dimer-sized lattice gap in 96 nm repeat structure from SS *doublet withODAs*. The gap is flanked by two additional densities (indicated by black arrowheads). The gap is filled in the *Mature* axoneme, and density at ‘Site 2’ is absent. D) Surface view of the A-tubule outer surface showing presence of the lattice gap in SS *doublet noODAs* and Rg *doublet withODAs* averages along with sequential addition of 96 nm repeat complexes. Position of RSP bases (1, 2 and 3) are highlighted.

Surprisingly, our subtomogram averages revealed tip-specific lattice gaps in the DMT (Figure 4B-D). The gaps correspond to the size of a tubulin dimer and recur every 96 nm in protofilament A02. Adjacent to these gaps, one copy of Rib72 and FAP252 and three copies of FAP115 are absent from the A-tubule lumen (Figure E4C), revealing the co-dependent binding of these MIPs and the missing tubulin dimer. Lattice gaps were present only in Stages 3 and 4, but not closer to the tip (Stages 1-2) or in the *mature* axoneme. This suggests that a tubulin dimer is actively and transiently removed during assembly and subsequently replaced.

We found additional densities on the outer A-tubule surface flanking each lattice gap (Figure 4C). One density is tip-specific whereas the other is also present in the axoneme shaft, although it was not reported in previous work ^3,4,31^. Since both densities contact FAP59/172 (mammalian CCDC39/40), we hypothesize that these are positional markers that coordinate tubulin removal and reinsertion at this site. All DMT MIPs are small enough to pass through the gaps (Figure E4D), so transient lattice openings could grant MIPs access to their luminal binding sites. This would overcome the spatial constraints imposed by the dense plug structures at the A-tubule ends and facilitate axoneme assembly.

### Stages 3-5: Sequential addition of motors and regulatory complexes to the DMT

Once the DMT architecture is established, large complexes required for ciliary beating assemble on its surface during Stages 3-5. In a mature axoneme, the ODAs, the main force-generating motors, bind every 24 nm via the ODA docking complex (ODA-DC), while the 96-nm repeating components, critical for regulation of ciliary waveform, bind to a separate surface of the DMT. Each 96 nm repeat contains three radial spoke (RSP) complexes, six single-headed inner dynein arms (IDAa-e, g), one double-headed dynein (IDAf), and the Nexin-dynein regulatory complex (N-DRC). Positioning of these elements is guided by the ccMAP FAP59/172 and stabilized through an interconnected network of docking factors. However, how this network is established during axonemal assembly to ensure proper beat generation is not known.

We found that RSPs and IDAs are sequentially recruited to the lattice in distinct blocks. FAP59/172 binds early in Stage 2 (Figure 2C) followed in Stage 3 by a highly interconnected functional block of RS2, RS3-stump (RS3S), N-DRC and most of IDAs (IDAc/e/d/g/f) (Figure 4D, (E5A). IDAe showed variable occupancy across conditions (Figure E5A) and so appears peripheral to this core block. The double-headed dynein IDAf first appears as a weak density in Stage 3 and becomes fully docked only in Stage 4, likely stabilized by contacts with the ODAs, which assemble at the same stage (Figure E5B). RS1 is recruited independently from the RS2/3S block, also in Stage 4 (Figure 4D, E5A), together with its tightly associated motor IDAa, forming a discrete RS1/IDAa unit. The last large complex to bind is IDAb, which only assembles outside the tip, in Stage 5.

Our results show that FAP59/172 is present on the lattice well before large 96-nm repeating complexes assemble, suggesting that their subsequent addition is actively regulated, possibly through controlled delivery or modification of the complexes and their docking factors. In contrast, ODAs bind concurrently with the components that facilitate their attachment, the ODA-DC and FAP182^6^. More broadly, the regulatory components (RSPs, IDAs and N-DRC) are recruited before the main force generators ODAs. This would prevent unregulated, premature force production during assembly and ensures structural reinforcement of the axonemal tip prior to ODA activation.

### Stage 5: Functional modularity is built into the assembly process

Axonemes have species- and cell type-specific structural adaptations that enable specialized functions. Consequently, some axonemal components are evolutionarily conserved, whereas others are species-specific. By mapping the assembly stage of *Chlamydomonas* MIPs and MAPs (Figure 5A, per-stage component list provided in Supplementary Information) to their conservation via phylogenetic orthogrouping (Figure 5B), we found that the order of assembly largely mirrors their evolutionary conservation. Notably, all MIPs added outside the tip, in Stage 5, are Chlamydomonas-specific, including FAP363, FAP166 and Rib30, which bind the wall between A- and B-tubules (Figure E6A). In mammals, this site instead contains Tektin filaments that reinforce the DMT ^6,9^. Similarly, we noted the absence of the B-tubule fMIP FAP112 from the tip (Figure E6B). The role of these Chlamydomonas-specific MIPs is currently obscure but this work shows they are dispensable for DMT formation. Species-specific components are thus incorporated at the final assembly step, potentially providing modularity to DMT formation by allowing the conserved DMT core to assemble first, followed by lineage specific factors that fine-tune axonemal mechanics.

**Figure 5.**
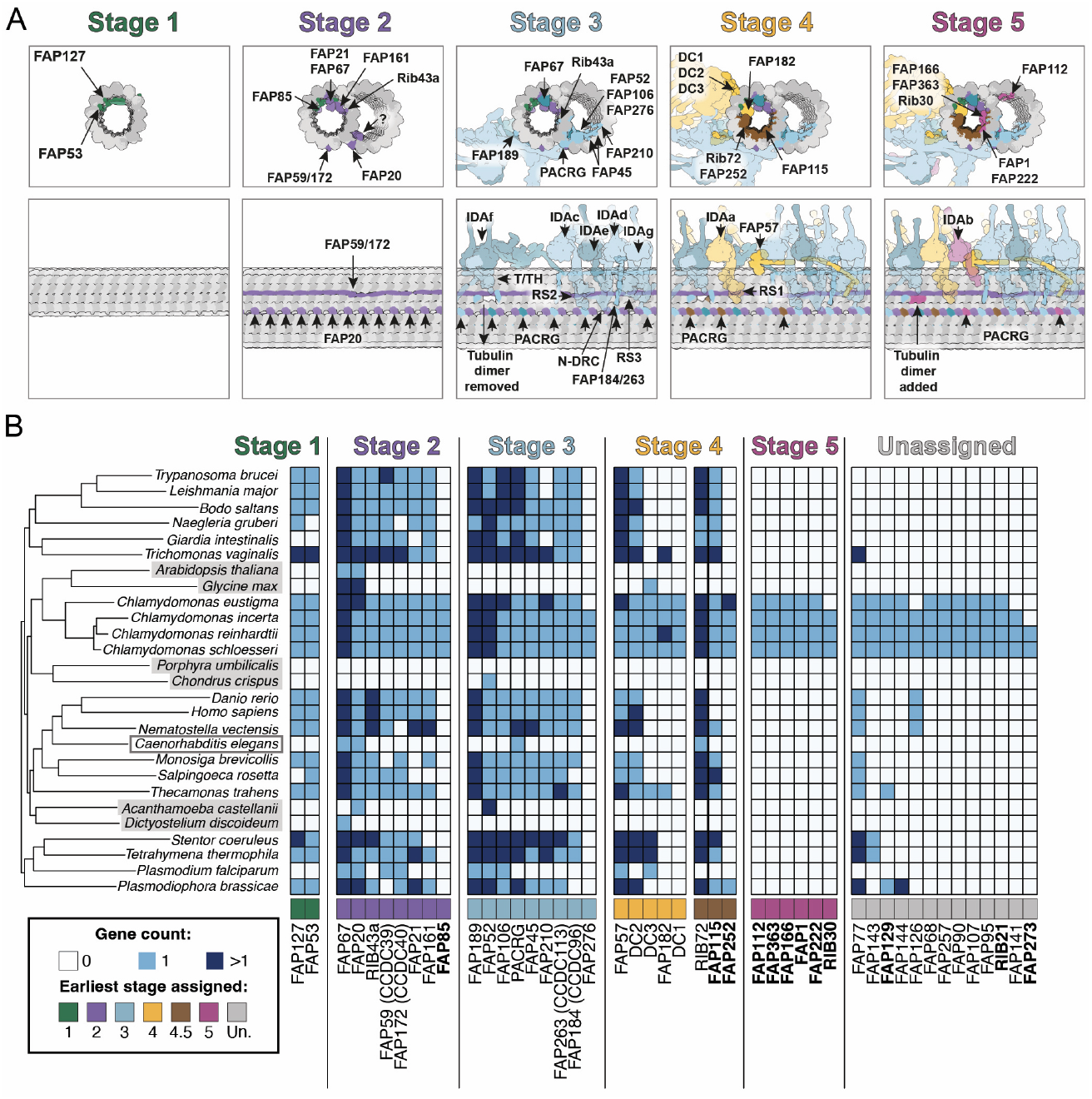
DMT assembly stages reflect phylogenetic conservation. A) Proposed model for DMT assembly, generated by integrating data from Rg and SS tips. Five stages are defined: Stage 1 (green) sees binding of the pathfinder fMIPs; Stage 2 (purple) sees incorporation of seam-binders and initial B-tubule scaffolding; Stage 3 (blue) marks IJ maturation and initial addition of large complexes to the outer DMT surface. Stage 4 (yellow) sees addition of ODAs and coating of the A-tubule lumen. Variably incorporated components are shown in dark blue (Stage 3.5) and brown (Stage 4.5). Stage 5 (pink) occurs outside the tip subcompartment. See Supplementary Information for full list of components present in each stage and evidence behind model generation. B) Table showing phylogenetic orthology groups for *C. reinhardtii* MIPs and MAPs, ordered with respect to their stage of incorporation. Where proteins are added in more than one stage, they are listed in the earliest stage. The number of orthologous genes identified is shown in blue. Species highlighted in grey lack cilia while *C. elegans* (grey box) only generate nonmotile cilia. Components highlighted in bold additionally lack structural homologues in mammals, based on experimental structures of motile cilia ^8^.

### Components in the C2 arm pioneer Central Apparatus formation

We next focused on the central apparatus CA, which is essential for the efficient beating of motile cilia and, like DMTs, is assembled at the growing tip. The CA consists of a pair of singlet microtubules (C1 and C2) which host four major projection arms (C1a, C1b, C2a, C2b). FAP20 and calmodulin are the only shared binding partners between the CA and DMTs ^1,2,4^. Unlike DMTs, the CA lacks a direct connection to the basal body structure ^12,13,48^. To understand CA assembly, we initially generated 8 nm averages of all four visible zones: LLS, *Bare, Zone 1, Zone 2* (Figure 1E-F). For *Zone 1* and *Zone 2*, we additionally generated 16 and 32 nm repeat averages of C1 and C2 (see Supplementary Information) and compared these in SS tips and *mature* axonemes.

The CP microtubule plus-ends are bound by the LLS^22^ and capped by the central microtubule cap (CMC), which is thought to regulate polymerization in a manner analogous to A-tubule plugs ^47^. Our in situ cryo-ET revealed that the CMC comprises three large rings in addition to short plug-like densities (Figure 6A-B) which are similar to the A-tubule plus-end plugs (Figure 4A, E1A). The first ring is flat and rigidly coupled to the LLS (Figure 6C), the third ring shows 9-fold symmetry and is membrane associated (Figure 6D). The two are tethered via the flexible intermediate (second) ring (Figure E7A-C). Adjacent to the third ring, on the extracellular side of the membrane, we observed a two-layered net-like structure with regular fibril spacing (Figure 6E-F, E7D) which is likely glycoprotein-based ^49^. We rapidly demembranated cilia and found the CMC and net remain physically coupled after this treatment (Figure E7E), suggesting the CMC links the LLS to the CP microtubule ends while spanning intra- and extra-cellular environments.

**Figure 6.**
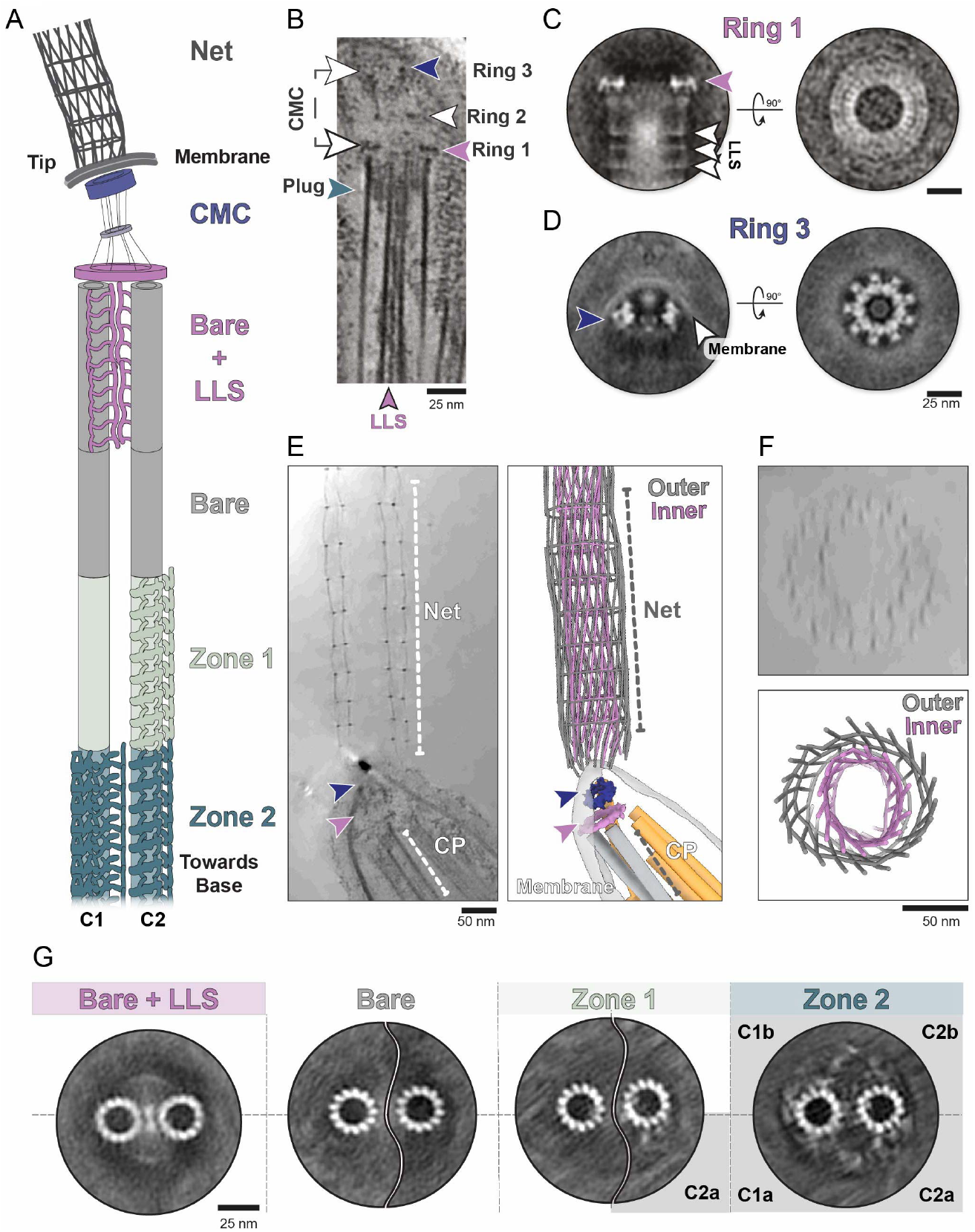
The CMC links the CP microtubules to an extracellular net. A) Schematic overview of CP morphology in the ciliary tip. B) Tomogram slice showing the CMC is composed of three rings and both CP microtubules contain short plug-like structures. C) Subtomogram average of Ring 1, showing it is a flat disc which sits just above the CP microtubules. The LLS is visible in the Ring 1 structure, showing they are in fixed position relative to each other. D) Subtomogram averaging of Ring 3, which exhibits 9-fold symmetry and membrane association. E) Tomogram (left) and 3D model (right) showing a double-layered extracellular net lies at the ciliary tip. F) Cross-section of the net from (E), showing the two concentric layers of the net (inner layer is pink, outer layer is grey). G) 8 nm repeat subtomogram averages of CP microtubule sections within SS ciliary tips. *Zone 1* features C2 decoration while *Zone 2* contains all four major CA projection arms.

Our subtomogram averages showed that the LLS is flexible and only loosely attached to the CP microtubules (Figure 6G). The zone directly below the LLS lacks MIPs and MAPs, so we call this the *bare* zone (Figure 6G). CA assembly instead begins in *Zone 1*, where the C2 microtubule gains decoration (Figure 6G). Here, the 16-nm repeating C2a binds along with the 8-nm repeating MAPs PF20 and FAP178 (Figure 7A-B). PF20 and FAP178 both localize at the seam and have been proposed to establish C2 periodicity ^1,2^. Consistent with this, mutation of PF20 leads to complete CA loss rather than partial disruption ^50^ supporting their foundational role in setting up the periodicity of the entire CA.

**Figure 7.**
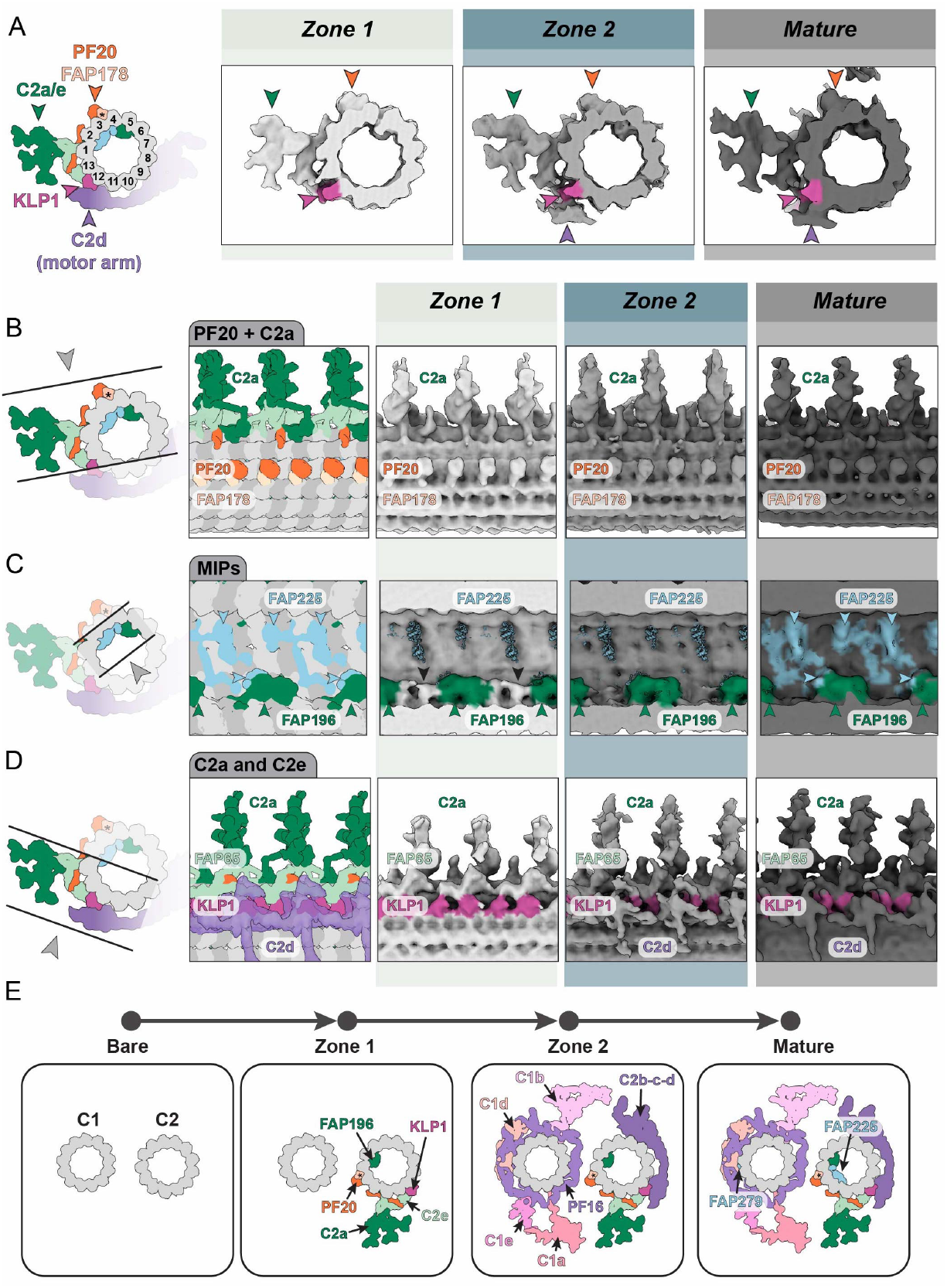
C2a components and KLP1 pioneer CA assembly. A-D) Panels showing comparison of 16 nm subtomogram averages of decorations on C2 microtubules in SS tips and the *mature* axoneme. Viewing perspective and components expected are highlighted in cartoons. A) C2 seam-binding proteins (FAP178/PF20), C2a/C2e arms and KLP1 bind in *Zone 1*, followed by the motor arm in *Zone 2*. B) Appearance of seam-binding components is consistent from *Zone 1* onwards. C) Progressive addition of FAP196 in the C2 lumen in *Zone 1* followed by FAP225 in the *mature* axoneme. Additional luminal density (black arrowheads) of unknown composition is present only in *Zone 1*. D) The KLP1 array and FAP65 are present in *Zone 1* while the C2d (motor arm) appears in *Zone 2*. E) CA assembly proceeds in two major steps. The C2a arm and seam-binding components bind first, in *Zone 1* and all major C1 projections and C2b-c-d are added in *Zone 2*. Two components are added outside the tip: FAP225 in the C2 lumen and FAP279 on the C1 surface.

Concurrently, the C2 lumen gains FAP196 and an additional unidentified MIP that binds only transiently, only in *Zone 1* (Figure 7C). On the C2 outer surface, C2e (containing FAP65) and, surprisingly, the KLP1 array are present (Figure 7D). KLP1 (mammalian Kif9) contributes to ciliary beat regulation by modulating the position of the ‘motor arm’ which wraps around C2 to contact C1^2,51^. KLP1 is the only ‘motor arm’ component in Zone 1, raising the question of how it is recruited. We performed an AlphaFold-based *in silico* screen and identified PF20 as the sole high-confidence interactor, with the flexible N-terminus of PF20 predicted to bind a helical hairpin in the KLP1 C-terminus (Figure E8A-D). Thus, KLP1 is loaded independently of other motor arm components and may be recruited to the lattice by PF20.

Near-complete CA assembly is achieved in *Zone 2*. At this stage, the remaining motor arm components (C2b-c-d) are added to C2 (Figure 6G, 7D), while C1, which was undecorated in *Zone 1*, becomes coated by C1a-e arms (Figure 6G, E9A). The simultaneous addition of C1a arms suggests that these large complexes are preassembled before integration and that their positioning is templated by C2 components. Compared to the *mature* axoneme, the CA in *Zone 2* lacks the EF-hand containing protein FAP225 in the C2 lumen (Figure 7C) and shows reduced FAP279 density in the C1d arm (Figure E9B). Thus, like DMTs, CA assembly continues beyond the distal tip into the axoneme shaft.

Overall, we found that CA assembly proceeds in two steps (Figure 7E). First, the C2a arm, PF20, FAP178, FAP65, KLP1 and FAP196 are added to bare microtubules. Next, all C1 projection arms are added simultaneously with C2b-c-d. The late addition of C2b-c-d suggests stronger tethering to C1 rather than C2, consistent with mutations in CPC1 or Hydin that remove C2b ^52,53^. Finally, our data show that the C1 and C2 microtubules stabilize their relative position only in *Zone 2*, after motor arm addition (Figure 6G, Supplementary Information), highlighting its role in fixing CP geometry.

## Discussion

In this study, we explain the assembly process of motile cilia axonemes by directly visualizing intermediate assembly steps at the ciliary tip using cryo-electron tomography.

DMT periodicity is thought to be established by MAPs, such as FAP57/172 (mammalian CCD39/40), that associate end-on-end, generating binding sites for other axonemal components with fixed spacing ^3,4,32^. Here, we show that the fMIPs FAP53 and FAP127 (CCDC11 and MNS1 in mammals) are the pathfinder components that establish DMT periodicity at the tip (Figure 2), meaning that the axonemal periodicity arises within the microtubule lumen of the singlet rather than through the external MAPs. Both FAP53 and FAP127 are present in the A-tubule in the basal body ^54^, suggesting their end-to-end self-association may propagate from base to tip alongside DMT extension. FAP53 and FAP127 are found in all studied DMTs ^3–6,8,9,44,45,55–58^ and in humans, mutations are associated with severe laterality defects ^59–64^, supporting a fundamental role in ciliogenesis. Mice lacking CCDC11 or MNS1 show tissue-specific phenotypic variation ^65,66^, suggesting partial redundancy in their activities.

The next stage in DMT assembly is the nucleation of the B-tubule at the outer junction OJ. The OJ is directly adjacent to the A-tubule seam, where the pathfinder fMIPs extend an α-helical arm through the microtubule lattice ^3^. Additionally, we propose that seam-binders including FAP67 (mammalian NME7), FAP161 and FAP85 are key drivers of B-tubule nucleation as their binding locally induces high curvature of the A-tubule (Figure 2, E2F). This could function to expose more of the pathfinders FAP53 and FAP127 on the external surface of the A-tubule, thus generating a defined B-tubule nucleation site. Removal of the tubulin C-terminus tails can promote formation of B-tubule-like structures *in vitro* ^67^, suggesting that the pathfinder MIPs could promote B-tubule formation by masking C-terminal tails at the OJ.

We found that B-tubule closure at the inner junction (IJ) occurs gradually, across different stages of the assembly process (Figure 3). Initially, a non-canonical intermediate transiently scaffolds B-tubule closure before being replaced by the canonical IJ components. Further work will be required to reveal the nature of the proteins which make up this transient and likely unstable non-canonical assembly intermediate.

In the mature IJ, FAP20 alternates with PACRG to form the “IJ filament”, but PACRG is initially absent from the earliest stage of IJ assembly while FAP20 is present. This shows that FAP20 and PACRG are incorporated independently and play distinct roles, a separation that was not previously evident from analysis of Chlamydomonas mutants^38^. Our conclusion is consistent with data from other organisms: in *C. elegans* sensory cilia only FAP20 is used at the IJ^5^, and in Trypanosomes, PACRG is dispensable for IJ formation if FAP20 is present ^68,69^. Our data shows that PACRG is progressively added, with alternating subunits appearing in Stage 3 and generating a 16 nm repeat. We observed staggered addition of apparently equivalent proteins also for FAP67 and Rib43a, showing this is a common theme in DMT assembly and suggesting their binding sites are progressively created, possibly by sequential addition of neighboring MIPs or MAPs or through accumulation of post translational modifications.

Further evidence for our model of regulated DMT formation is the transient A-tubule lattice gaps we observe at the tip (Figure 4). In our averages (Figure 5), A-tubules initially form without gaps (Stages 1-2), after which a single tubulin dimer is removed at a fixed position in the 96nm repeat (Stages 3-4) and later filled again in the axoneme shaft (Stage 5). Because tubulin normally forms complete microtubules and the energy required to remove a tubulin dimer from of the lattice side wall is ∼30pN ^70^, this removal cannot happen without an active process. We also detect additional, unassigned protein densities flanking the gap, which we propose regulate its formation and re-closure. Since many luminal A-tubule MIPs are added far from the tip and the A-tubule plus end is blocked by plugs, lattice gaps can provide essential access for late-added MIPs to reach their binding sites.

We propose that functional modularity is an inbuilt feature of axoneme assembly. For example, minor IDA isoforms are expressed during early stages of ciliogenesis ^46,71,72^ but their structure, function, and precise distribution remain a mystery. Our data shows IDAb and IDAe are variably added during assembly and so these may be more easily replaced than the other IDAs for adaptation of the 96 nm repeat architecture.

Similarly, the RSPs undergo staggered recruitment, suggesting functional differentiation. RSPs are essential for proper mechanoregulation of the ciliary beat, yet the contribution of individual RSPs is not well understood. Most RS1 and RS2 proteins are shared, with only 3-4 unique components per complex ^73,74^ The major docking factor for RS2 is FAP9^73^, an elongated helical protein which threads through the N-DRC to form a key part of RS3S. miRNA knockdown of FAP91 reduces both RS2 and RS3S components ^75^, supporting our observation that they are recruited together. In contrast, RS1 docking is mediated by FAP253 and our data shows that the RS1 binding site is generated after that of RS2/3S, simultaneously with the ODAs. This staggered recruitment supports functional differentiation between RS1 and RS2.

The CA provides essential mechanochemical regulation of the ciliary beat and we find it assembles sequentially, similar to DMTs. We identified two major steps in CA assembly and a third occurring beyond the ciliary tip. Unlike DMTs, where MIPs extend to near the microtubule plus ends, the CP microtubules appear undecorated over their distal ∼400 nm (Figure 1E-F, 6G). Periodicity is first established in *Zone 1* through partial C2 decoration, followed by concurrent addition of the remaining C2 and C1 projection arms in *Zone 2*. The simultaneous incorporation of large CA complexes suggests preassembly before ciliary import, as seen for DMT components like RSPs^76,77^ and ODAs^78^. This would streamline tip assembly and enable rapid axoneme formation.

Tight coupling of the C1 and C2 microtubules and interlinkage of CA projections along the axonemal shaft ^79^ is essential for motile cilia function, as mutants with unstable C1-C2 alignment retain both microtubules but display severe defects in beat coordination. Consistent with this, our finding that stable C1-C2 positioning emerges only late in assembly together with motor arm incorporation, suggests that mechanical stabilization of the central apparatus and regulatory competence are established simultaneously.

How C1-C2 alignment is achieved remains a central question. A seam-centric model ^1,2^ proposes that seam-binding components define the periodicity and orient the central pair microtubules. Consistent with this, we found that the C2 seam binders FAP178 and PF20 associate early in *Zone 1*, providing a positional reference that can subsequently be read by the C1 components FAP47 and Hydin, which mediate flexible C1 and C2 interactions ^1,2^. Hydin, also part of the C2b-c-d motor arm, may also integrate positional cues by reading KLP1/FAP65 periodicity on C2 to further align C1 and C2 at the seam.

Our data further refine the role of KLP1 in CA assembly and function. KLP1 is essential for ciliary motility ^80,81^ and was thought to be inseparably linked to the motor arm, as its loss leads to C2b absence ^81^ and its conformation dictates motor arm positioning ^2^. However, we our data show that *Zone 1* components are sufficient to recruit KLP1 to the CA, independent from the motor arm. Mutations in PF20 and FAP65, both adjacent to KLP1, severely disrupt CA assembly ^50,82–84^, suggesting that they aid KLP1 recruitment in *Zone 1*. This is consistent with our *in silico* prediction that PF20 directly interacts with KLP1 (Figure E8), and underscores the fundamental role of PF20 in guiding KLP1 incorporation and CA assembly. Our data additionally show the C2b-c-d motor arm assembles separately from the other C2 components and KLP1, meaning these are not constitutively coupled and suggests that motor arm slippage is tolerated, with KLP1 serving as a structural guide for central pair organization rather than a binary switch for the motor arm.

Overall, our in situ cryo-ET analysis shows that axoneme assembly proceeds through defined structural intermediates and extends well beyond the ciliary tip. DMTs and the CA are constructed through sequential incorporation of functionally distinct modules, with periodicity, mechanical coupling, and motor activity established in a regulated manner. This assembly logic ensures coordinated emergence of structural integrity and motility while allowing flexibility for species- and cell type-specific adaptation. Thus, our work provides a blueprint for understanding how disruptions at specific stages of assembly lead to axonemal dysfunction.

## ACKNOWLEDGEMENTS

We would like to thank P. Swuec and S. Sorrentino (Cryo-electron Microscopy Unit, National Facility for Structural Biology, Human Technopole), A. Graziadei (Structural Proteomics Unit, National Facility for Structural Biology, Human Technopole), D. Dalle Nogare and E. Cammarota (Bioimage Analysis Infrastructure Unit, National Facility for Data Handling and Analysis, Human Technopole), C. Fernandez and D. Colombo (IT Department, Human Technopole) for technical support. We acknowledge Mareike Jordan for sharing preliminary observations which are described in her PhD thesis (https://katalog.slub-dresden.de/en/id/0-1760015911). We acknowledge funding from Human Technopole, the European Research Council under the EU Horizon 2020 Research and Innovation Programme (grant number 819826) to G.P, an GAČR-DFG Cooperation Grant (#665136) to G.P., and an EMBO Long Term Fellowship (ALTF 1141-2021) to H.E.F.

## RESOURCE AVAILABILITY

Structural data supporting the findings are available via the EMDB and their analysis detailed in the Supplementary Information. Doublet 8 nm repeat structures are EMD-XXXXX to EMD-XXXXX; Doublet 96 nm repeat structures are EMD-XXXXX to EMD-XXXXX; C1 32 nm repeat structures are EMD-XXXXX and EMD-XXXXX; C2 16 nm repeat structures are EMD-XXXXX to EMD-XXXXX; Microtubule capping structures are EMD-XXXXX to EMD-XXXXX. Any additional materials are available from the corresponding authors upon request.

## DESCRIPTION OF SUPPLEMENTARY INFORMATION

Supplementary Information contains Figures 1-14 show subtomogram averaging and classification methodologies and summary of evidence used to generate models of assembly for the DMT and CA. Supplementary Movies 1 and 2 show molecular animations of the DMT and CA assembly processes described in this work. Supplementary Table 1 contains genomes used for orthology inference for DMT MIPs and MAPs. Supplementary Table 2 contains list of Chlamydomonas proteins selected for orthology inference. Supplementary Table 3 contains orthologous proteins identified.

## AUTHOR CONTRIBUTIONS

Conceptualization, H.E.F. and G.P.; Writing - Original draft H.E.F. and S.E.L.; Writing - Review & Editing H.E.F, S.E.L. and G.P.; Investigation, H.E.F., S.E.L, and H.N.V.; Formal analysis. H.E.F, S.E.L., H.N.V., J.D-V., Visualization, H.E.F., S.E.L., M.R., Supervision, G.P., Funding acquisition, H.E.F. and G.P.

## Extended Data

**Figure E1.**
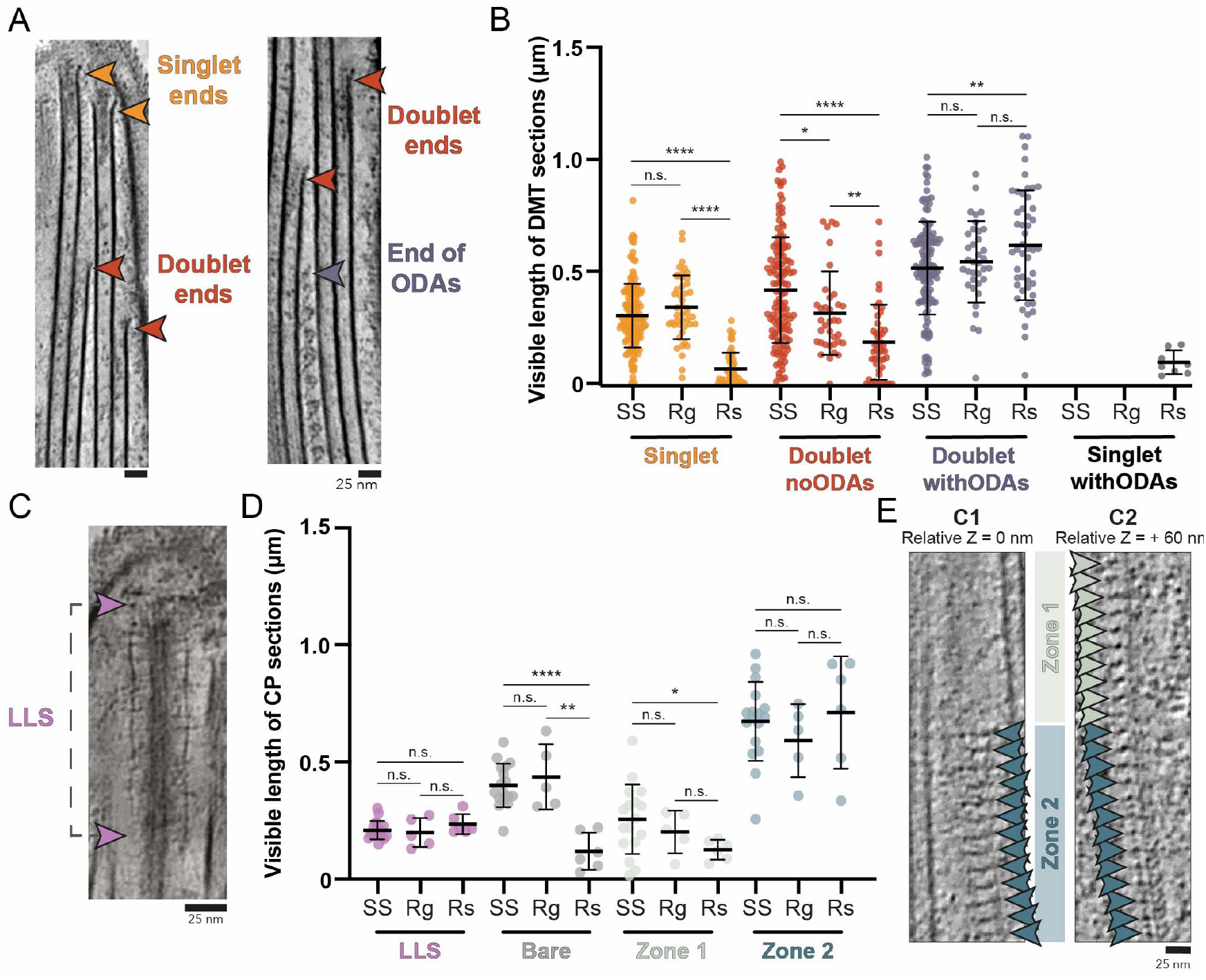
Quantitative comparison of DMT and CP architecture across growth states. (A) Tomogram slices showing landmarks used for DMT tracing. Singlet ends contain plugs while the B-tubule ends are open. B) Measured length of each DMT section measured across SS, Rg and Rs conditions. Mean ± SD and significance from pairwise two-tailed T-tests are shown. *doublet withODAs* lengths depend on tomogram field of view and are not shown in Fig 1D but included here for completeness. C) Example of LLS in denoised tomograms. The CP MTs are above and below this viewing plane. D) As in (B) but for CP. E) Repetitive densities on C1 and C2 microtubules used to distinguish *Zone 1*, (C2 only decorated) and *Zone 2* (both decorated).

**Figure E2.**
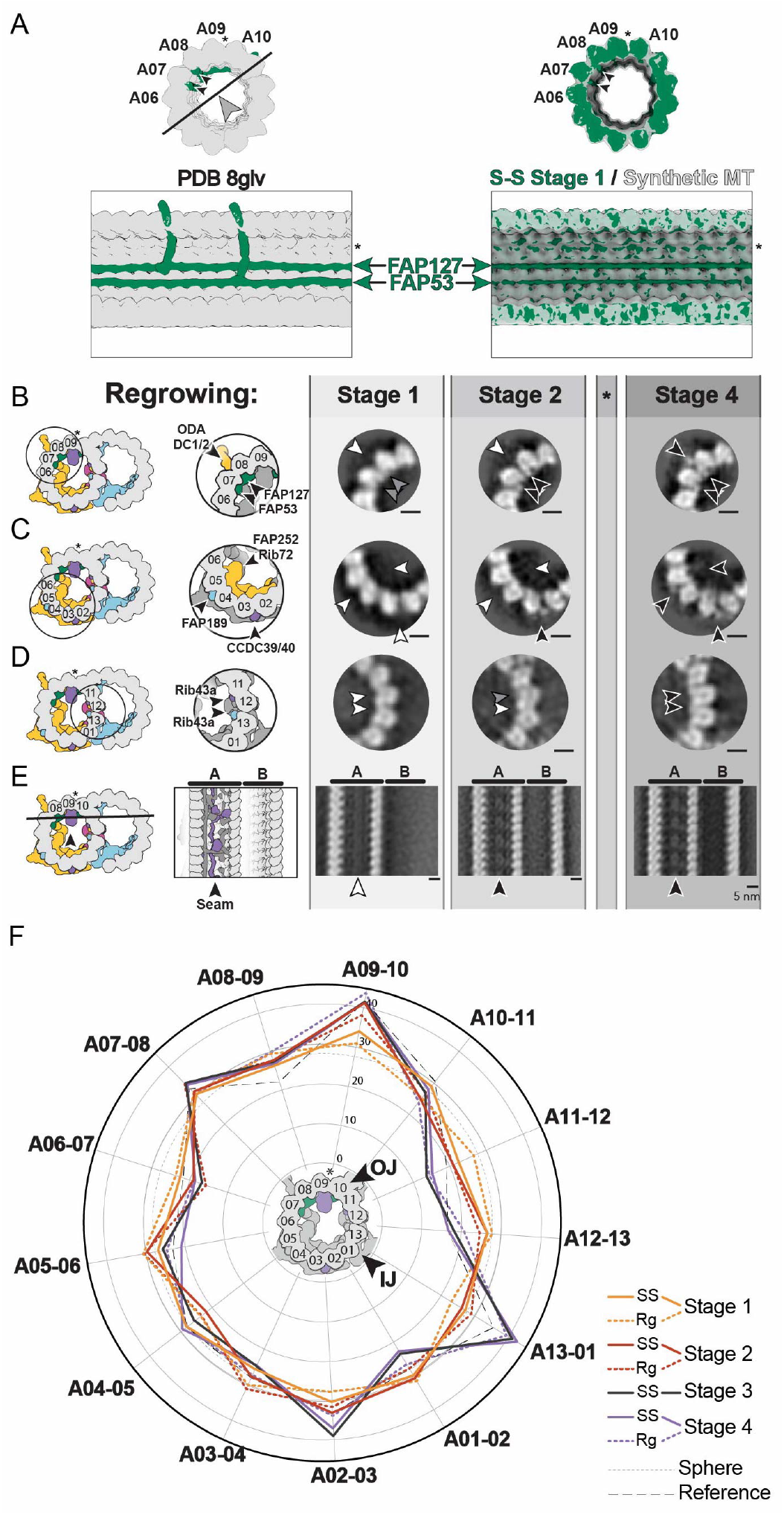
Both steady state and regrowing cilia tips undergo structural transitions during assembly. A) Tomogram slices showing landmarks used for DMT tracing. Singlet ends contain plugs while the B-tubule ends are open. B) Measured length of each DMT section measured across SS, Rg and Rs conditions. Mean ± SD and significance from pairwise two-tailed T-tests are shown. *doublet withODAs* lengths depend on tomogram field of view and are not shown in Fig 1D but included here for completeness. C) Example of LLS in denoised tomograms. The CP MTs are above and below this viewing plane. D) As in (B) but for CP. E) Repetitive densities on C1 and C2 microtubules used to distinguish *Zone 1*, (C2 only decorated) and *Zone 2* (both decorated).

**Figure E3.**
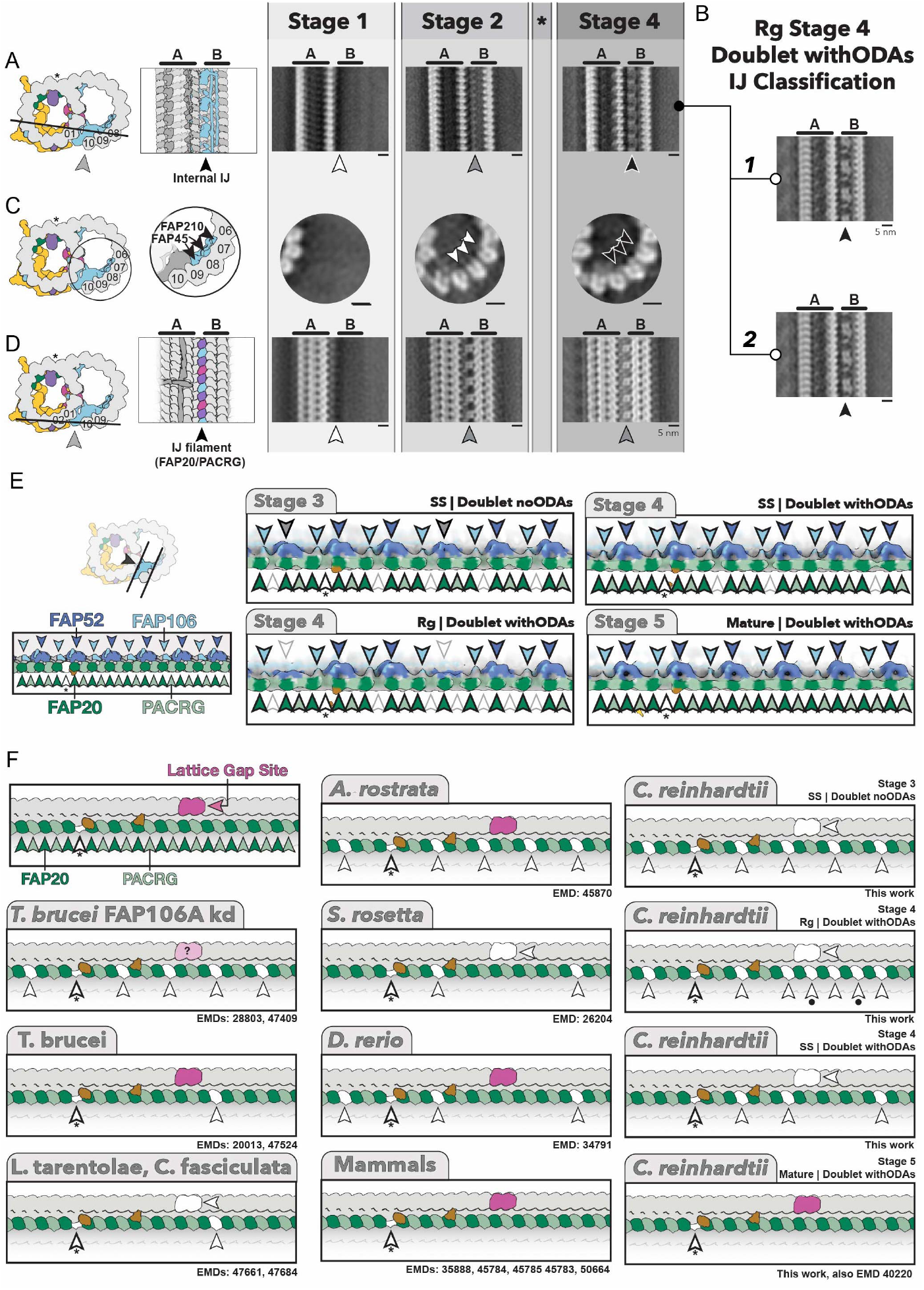
Partial IJ filament formation is conserved. A) Subtomogram average projections from each stage of DMT assembly showing appearance of inner IJ architecture in Rg cilia tip. B) Classification of *doublet withODAs* section in Rg tips, showing the expected 16 nm repeat architecture is obtained. C) As in A, showing binding of fMIPs FAP210 and FAP45. D) As in A, showing progressive IJ filament formation. E) Internal IJ components in the 96 nm repeat averages showing full occupancy apart from in the Rg Stage 4 average, where one copy of FAP52 is absent every 48 nm. Left cartoons show viewing perspective and components highlighted. F) Schematics showing PACRG occupancy across organisms. Position of expected components are labelled in top left panel. White arrowheads missing components. Subunits in the PACRG-A site are only absent in Rg *doublet withODAs* average (black dots), with the remainder in the PACRG-B site. Question mark indicates uncertainty in tubulin dimer status since this is based on a 48 nm repeat structure.

**Figure E4.**
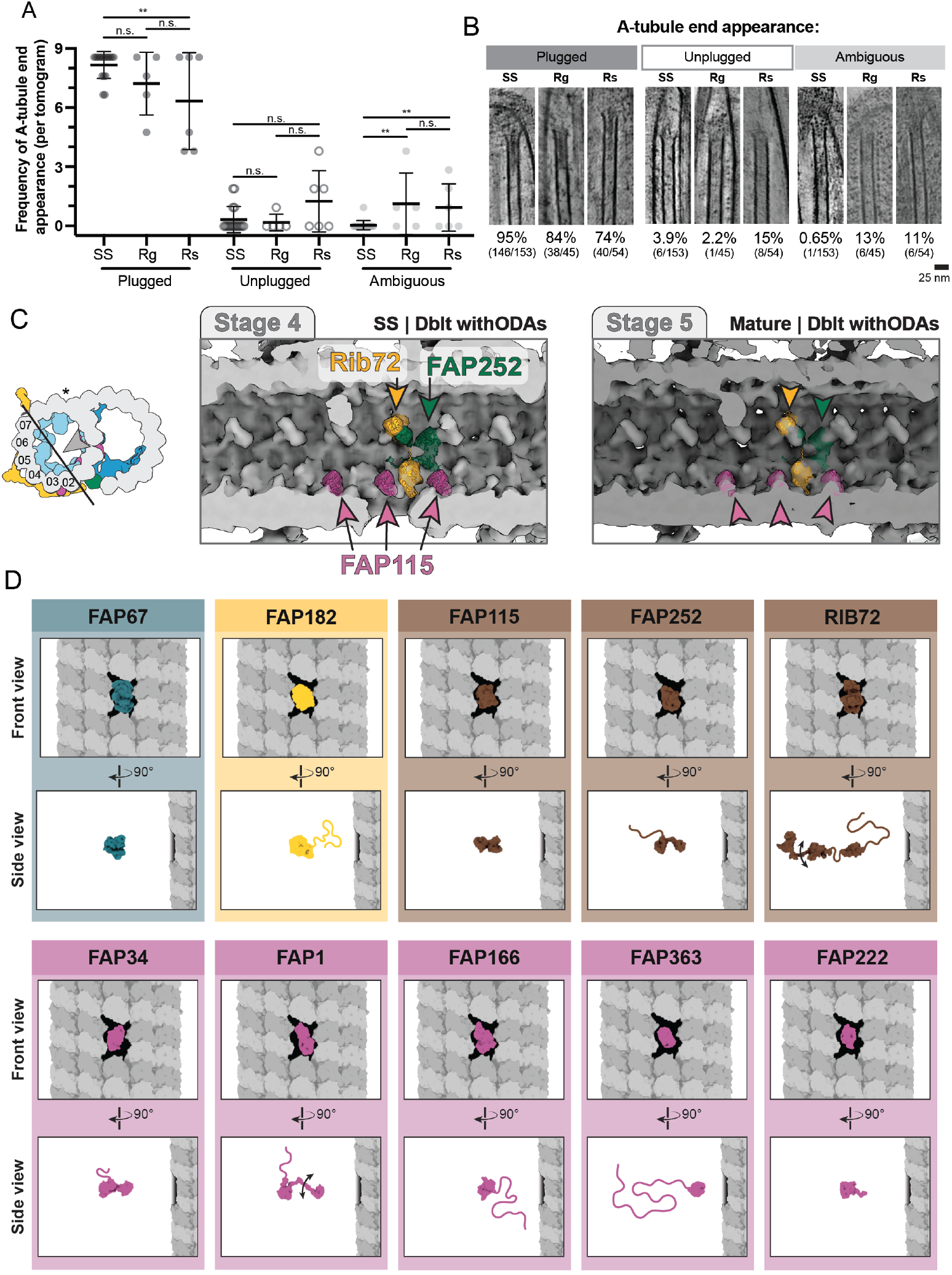
A-tubule components can gain luminal access through lattice gaps. A) Quantification of *singlet* A-tubule ends appearing plugged, unplugged or ambiguous in SS, Rg and Rs conditions. B) Slices through tomograms showing examples of singlet plugs for each category in (A). C) Cross-section through the A-tubule showing a subset of luminal MIPs are absent close to the lattice gap. Although usually repeating every 8 nm, one copy of Rib72, one copy of FAP252 and three copies of FAP115 are missing from the SS *doublet withODAs* 96 nm repeat average. D) 3D molecular modelling showing that MIPs can access the A-tubule lumen via lattice gaps. Front and side views are shown.

**Figure E5.**
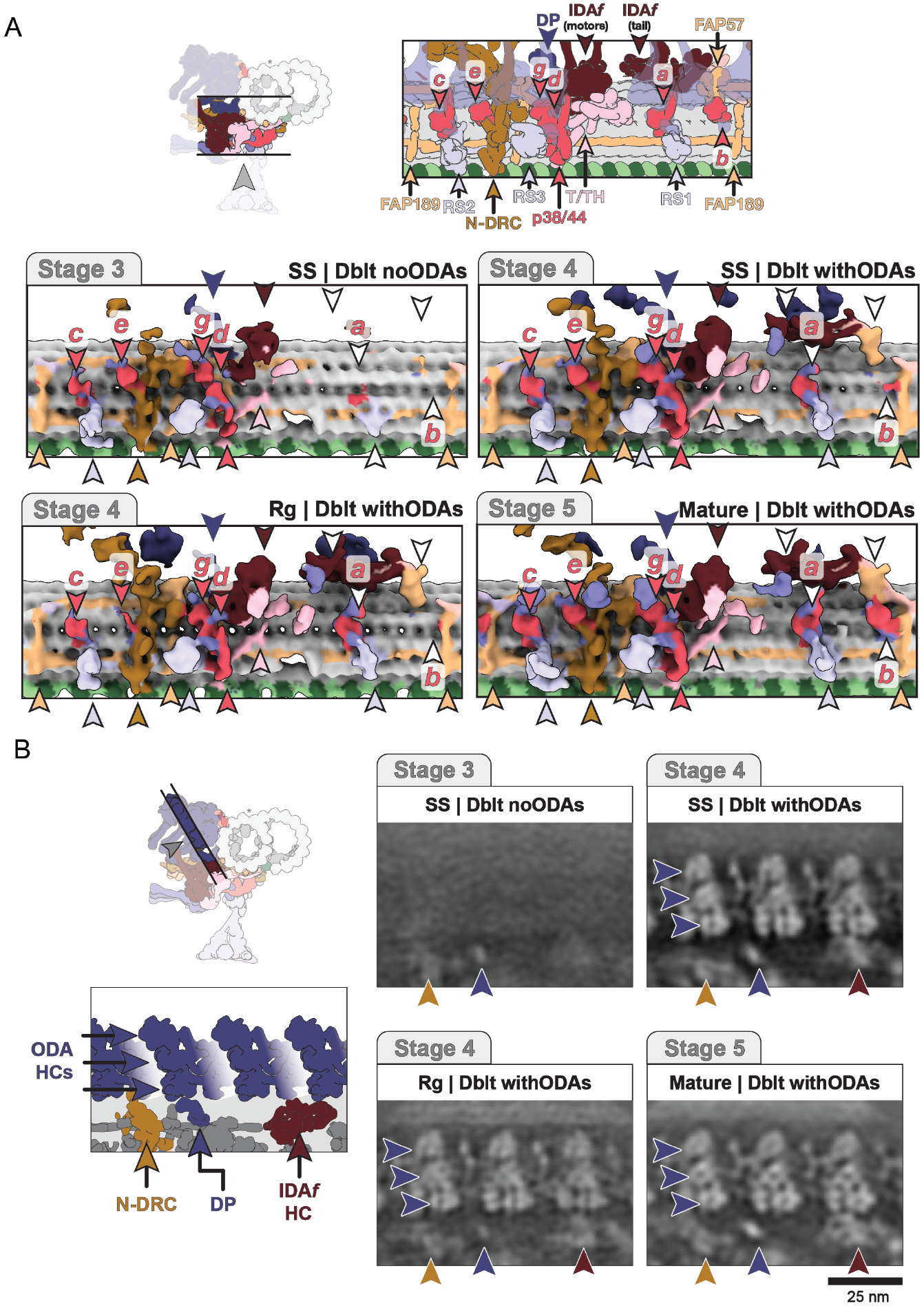
Binding of IDAs, ODAs and RSPs is sequential. A) Surface view of the A-tubule outer surface showing progressive addition of single-headed IDAs (IDAa,b,c,g,e purple) and double-headed dynein (IDAf, maroon), along with their docking factors (coral). B) Projections through 96 nm repeat averages showing that ODAs bind in Stage 4, after N-DRC addition.

**Figure E6.**
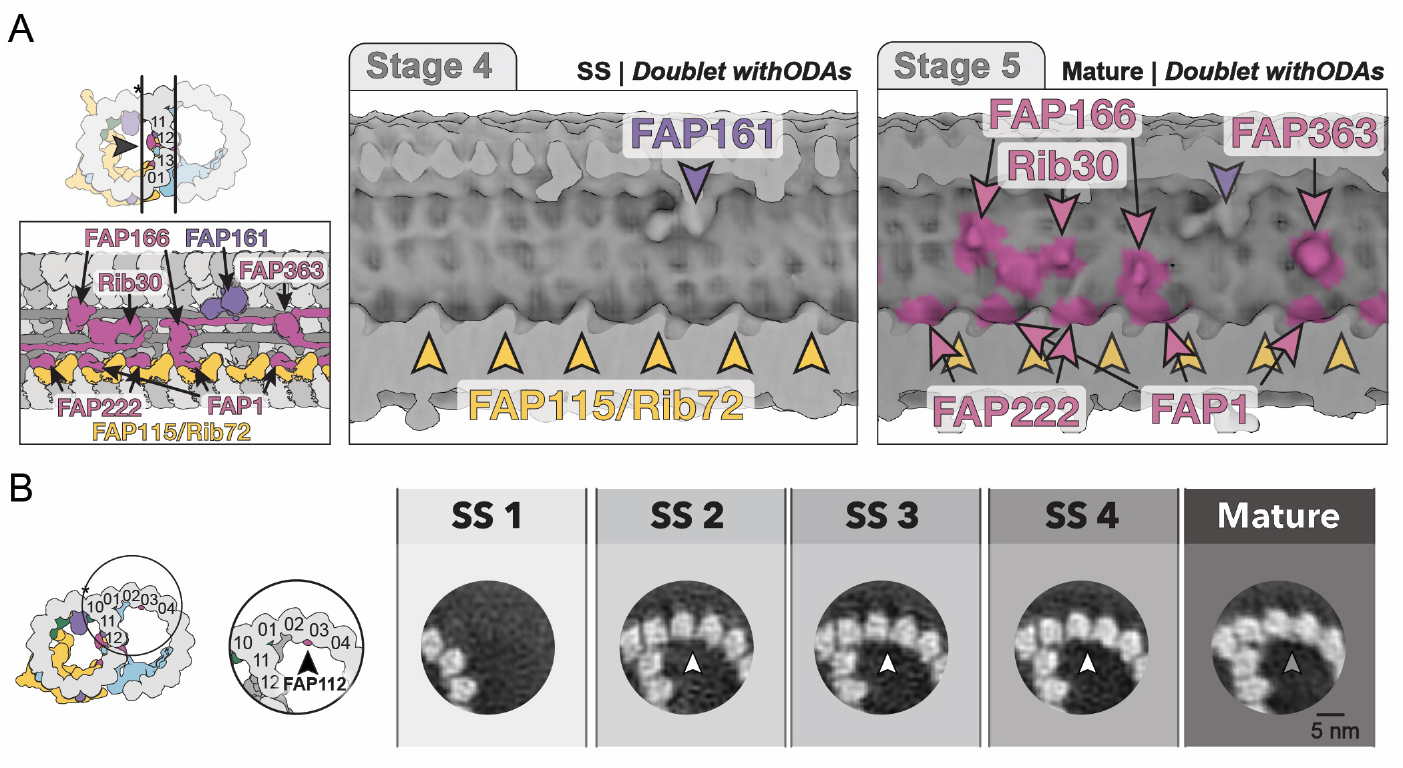
Chlamydomonas-specific MIPs are incorporated late in assembly. A) Surface view of the A-B tubule wall in 96 nm repeat showing Chlamydomonas-specific MIPs (FAP166, FAP363, Rib30, FAP1 and FAP222; pink) are absent from the tip. Viewing perspective and highlighted components are shown on the left. B) Projections of 8 nm repeat subtomogram averages showing FAP112 is absent from the tip.

**Figure E7.**
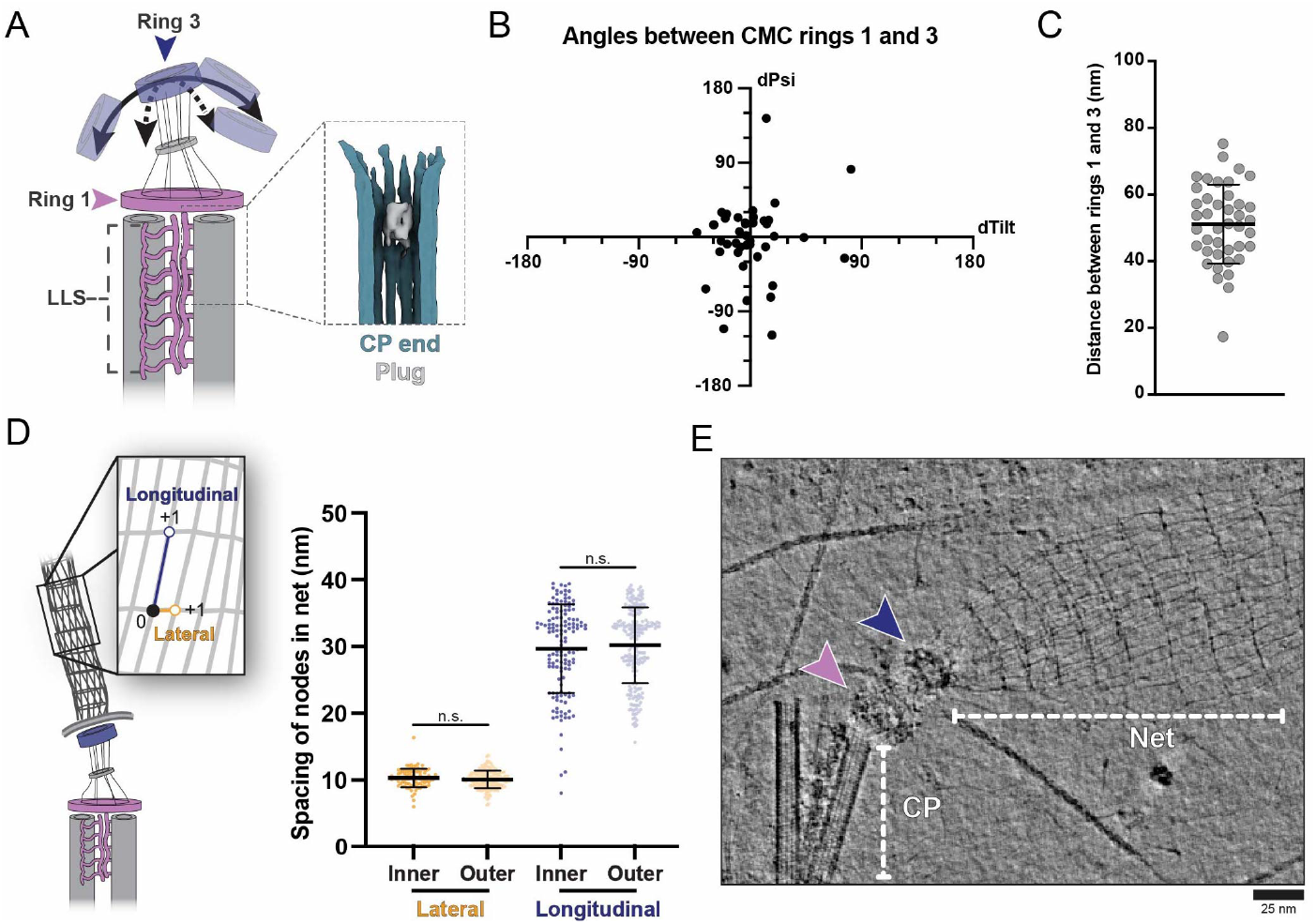
The CMC and net are flexibly tethered. A) Subtomogram average of CP ends shows short plugs are present in the lumen. B-C) High variability in angle (B) and distance (C) between Ring 1 and 3 in the CMC indicates they are loosely tethered. D) The two layers within the net have similar inter fibril spacing, as shown by measurement of the longitudinal and lateral spacing between nodes within the net. Mean ± SD and significance from pairwise two-tailed T-tests are shown. E) On-grid detergent extraction shows the CMC physically couples the net to the CP microtubule ends at the tip.

**Figure E8.**
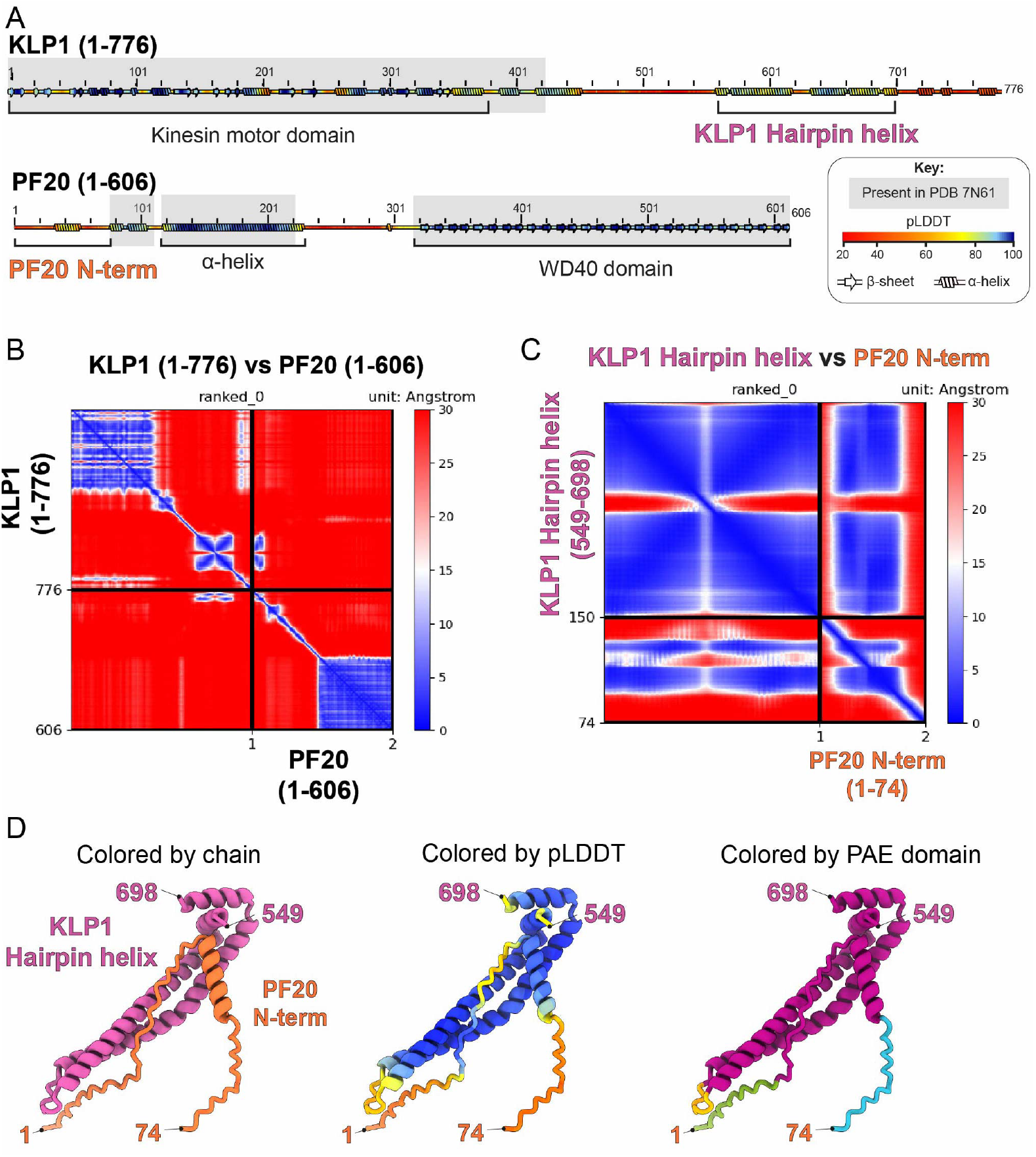
AlphaFold2 predicts PF20 as a KLP1 interaction partner. A) Ribbon plot of KLP1 and PF20 showing secondary structure based on AlphaFold2 predicted structures of individual components, colored by pLDDT. Regions present in PDB 7N61 are indicated along with the subregions predicted to interact. B) PAE plot for AlphaFold2 multimer predictions of full-length KLP1 and PF20. C) As in B), for subregions predicted to interact in initial screen. D) Cartoon representation of complex predicted to form between the flexible N-terminus of PF20 and a hairpin helix in KLP1.

**Figure E9.**
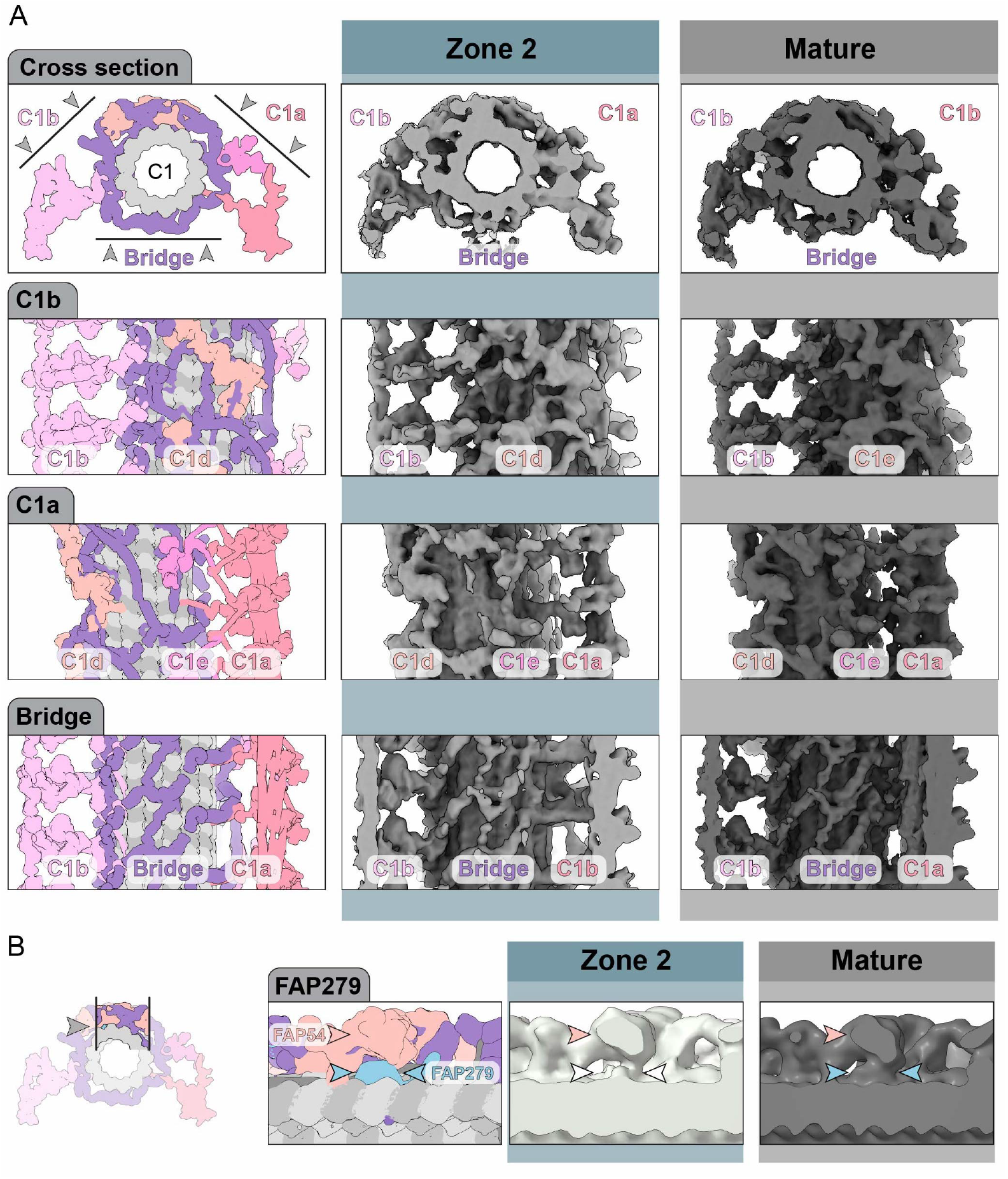
Major C1 projections assemble simultaneously in Zone 2. A) Panels showing comparison of C1 32 nm repeat subtomogram average structures in *Zone 2* and *mature* axoneme. Viewing perspective and projection arms in view are shown in cartoons. Overlay is shown to highlight the similarity in density observed across these assembly stages. B) Longitudinal cross section showing weak density for FAP279 in the Zone 2 C1 subtomogram average, suggesting it is added outside of the tip.

## Methods

### *Chlamydomonas reinhardtii* cell culture

Wild-type *Chlamydomonas reinhardtii* (strain 32M CC124) were cultured in Tris-acetate-phosphate (TAP) medium supplemented with Hutner’s trace elements obtained from the Chlamydomonas Resource Center (https://www.chlamycollection.org/product/hutners-trace-elements/). At least three days prior to experiments, cells were transferred from TAP-agar plates (0.1% agarose in TAP) into 300 mL liquid TAP medium and cultured in a 12-hour light/dark cycle at 22°C with constant aeration. To generate cells with regrowing (Rg) cilia, cells were subjected to pH shock. Briefly, 50 mL of culture was pelleted twice at 800 x g for 3 minutes and resuspended in 5 mL 10 mM HEPES (pH 7.5). Then, 75 µl 0.5 M Acetic acid was added, and after 30 s with gentle mixing, 37.5 µl 1M NaOH was added to restore the pH to 7.5. TAP medium was added to bring the final volume to 50 mL. Complete deciliation was confirmed by brightfield microscopy (EVOS M5000). Cilia regrowth was allowed to proceed for 20-40 minutes before cryo-EM sample preparation. To generate cells under resorbing (Rs) conditions, cells in TAP medium were treated with 10 mM or 20 mM sodium pyrophosphate (NaPPi). Ciliary resorption was allowed to proceed for 20 - 40 minutes before cryo-EM sample preparation.

### Sample preparation for cryo-EM

EM grids (Quantifoil R3.5/1 Au 200 mesh) were plasma cleaned for 10 s using a SOLARUS II Model 955 (Gatan) with an 80:20 oxygen/hydrogen gas mixture. Plunge freezing was performed using a Leica EM GP2 set at 22 °C and 95 % humidity by single-sided blotting. 3.5 µL of cell suspension was applied to the carbon side of the grid followed by 1.5 µL 10 nm BSA-coated gold fiducials (BBI solutions, diluted 1:20 in PBS). After blotting for 2s, grids were plunged into liquid ethane held at -184 °C, transferred to liquid nitrogen for storage and clipped into AutoGrids (Thermo Scientific) prior to imaging. For visualization of the connection between the extracellular net and CP microtubule, which is mediated by the CMC, we performed on-grid detergent extraction. Cells were prepared as described above and added to grids but instead of gold fiducials, 1 µL 0.005% NP-40 was added immediately prior to plunge freezing.

### Cryo-ET data acquisition

Tilt series were acquired on a Titan Krios G4 transmission electron microscope equipped with a Falcon 4i direct electron detector and Selectris X energy filter (Thermo Scientific). The microscope was operated at 300 keV with a slit width of 20 eV and nominal magnification of 33,000 x, to give pixel size of 3.72 Å and 1.5 µm field of view. Tilt series were acquired from + 60°to - 60°in 2°increments using a dose-symmetric tilt scheme ^85^. The target defocus was set between -3 and -5 µm and the dose per tilt was 1.9 e^-^/A^2^, resulting in a total dose of 120 e^-^/A^2^. Images were saved in EER file format and microscope control was performed using SerialEM ^86^.

### Tomogram reconstruction

Initial image processing and tomogram reconstruction were performed using the TOMOMAN pipeline ^87^. Raw tilt series were imported and motion-corrected MotionCor2 as implemented in RELION 3.1^88^, with frame grouping optimized to yield 5-7 sub-images per tilt for motion detection. Even/odd tilt images were saved for subsequent denoising. CTF estimation was performed using CTFFIND4^89^ and dose weighting applied using IMOD ^90^. After manual inspection to exclude poor-quality images (e.g. occluded fields of view or misaligned tilts), tilt series were aligned using gold fiducials in the eTomo package of IMOD. Next, bin4 (14.88 A/pix) tomograms were generated and filtered using a MATLAB implementation of WARP’s tomogram deconvolution filter ^91^, available at https://github.com/dtegunov/tom_deconv. Finally, a subset of tomograms were denoised using CryoCARE^92^, with networks being trained and applied for each tomogram individually. We selected high quality tomograms for analysis (16 from steady state, 5 each from regrowing and resorbing conditions) which were accurately aligned, had signal-to-noise ratio and cilia tips well supported by vitreous ice. Final tomograms were further deconvolution filtered for visualization of axonemal components at the ciliary tip.

### 3D modelling of tomograms

Denoised, filtered bin4 tomograms were manually inspected and modelled in IMOD. Open contours were drawn along the center of each microtubule, with points placed every 200 - 300 nm starting at the tip. For DMTs, the end of the A- and B-tubules, and distal-most ODA densities were marked. These positions were used to divide each contour into three sections and these were stored as separate objects within the IMOD model. For CP microtubules, the ends and the furthest extent of repeating densities on C1 and C2 were used to define the *bare* region, *Zone 1* (where C2 had decoration) and *Zone 2* (where both MTs had proteins bound). C2 was distinguished from C1 by visualization of repeating luminal densities. The length of the LLS was measured from the CP ends and overlapped with the *bare* region in SS and Rg conditions. In the Rs conditions, the *bare* region and *Zone 1* were shortened so the LLS often overlapped with the repetitive densities in *Zone 1* and *Zone 2*. MT lengths were measured and plots generated using imodinfo (an IMOD command). The lengths of the proximal-most section of each set of MTs (*Doublet withODAs* for DMTs, *Zone 2* for CP) depend on the tomogram field of view and so were not taken into consideration when comparing the architecture across conditions. A-tubule plug counts were also carried out using IMOD. We categorized any cases where the plug start or end points were unclear as ‘ambiguous. For extracellular net analysis, we measured the inter-fibril spacing by placing points at node crossovers and measuring the lateral or longitudinal distance to the next node using imodinfo.

### Initial subtomogram averaging of DMTs

The IMOD models described above were used for semi-automatic particle picking of DMTs. Supplementary Information contains details of DMT subtomogram averaging workflow and results. For each section (e.g. *Doublet withODAs*), the corresponding object was extracted into a new IMOD model using the ‘imodextract’ command. STOPGAP-compatible motivelists ^93^ were generated using custom MATLAB scripts incorporating functions from the TOM Toolbox (https://www.biochem.mpg.de/6348566/tom_e). During this process, the coordinates were super sampled every 4 nm, and Euler angles assigned based on the vector between adjacent particles, preserving the polarity from the IMOD model. Next, the STOPGAP motivelists was converted into RELION-compatible .star files using the dynamo2m function (https://github.com/alisterburt/dynamo2m). During this process, each microtubule was assigned a unique ID stored in the field ‘rlnHelicalTubeID’, to allow downstream unification of angles on a per-MT basis. Particles for each section (singlet, doublet noODAs, doublet withODAs) were extracted separately and assigned distinct rlnOpticsGroup numbers. Tilt series were imported into Warp, and subtomograms and 3D CTF particles extracted at bin2 (7.44 A/pix) using these .star files. The output .star files were then combined into a single star file for downstream processing. After subtomogram extraction, tilt series alignment was refined using M ^94^. For this, we performed 3D refinement of the pooled DMT subtomograms in RELION 3.1.3^88^. We retained the psi and angles derived from the MT orientation in the tomograms and randomized the rot angle before performing local 3D refinement with a cylindrical mask focused on the A-tubule. Local refinement in RELION was performed by setting the angular sampling and local search to the same value, typically 3.7°. Duplicate particles within 3 nm of each other were removed prior to tilt series refinement. Next, we re-extracted subtomograms at their original positions and the rot angle for each DMT was determined via 3D refinement. The previously refined tilt and psi angles were used as priors (rlnAngleTiltPrior, rlnAnglePsiPrior) with angular sampling 7.5°, local search 3.7° and additional arguments --sigma_tilt 3 and --sigma_psi 3 specified. *Singlet* subtomograms were excluded from this refinement step, as the goal was to determine the B-tubule orientation, which is absent in the singlet. Following this, median Euler angles were calculated and applied on a per-DMT basis using a Python script which uses the starfile package (https://github.com/teamtomo/starfile). These unified angles were applied to all particles from the corresponding DMT (including singlet) and a subsequent round of local 3D refinement was performed. This per-DMT unification of Euler angles was essential for resolving the fMIPs in the singlet section, since singlet lacks strong orientation cues, global refinement was insufficient. the signal for these components was too low to be latched onto in global refinements. By propagating orientation information from B-tubule containing sections we were able to resolve the *singlet* fMIPs. The star files were then cleaned to remove duplicate particles and the 8 nm repeat along each microtubule retained. Per-section subtomogram averages were then generated using a python-based script which uses the starparser package (https://github.com/sami-chaaban/starparser). One out of the nine DMTs in mature axonemes are expected to lack ODA along their entire length, as DMT1 characteristically contains a bridge-like structure that contacts the adjacent DMT ^71,95^. To reduce complexity for classification we excluded *doublet noODAs* subtomograms coming from DMTs completely lacking ODAs. These steps described until this point enabled generation of 8 nm repeat averages for the three DMT sections observed in denoised tomograms. Each DMT section was then masked and locally refined in up to 7 subregions (shown in Supplementary Figure S1). We generated references for each section from the pooled refinements above and used equivalent masks for all sections.

### IJ classification and stage definition

To analyze the IJ architecture, we performed classification within the A12-13/B09-10 region. In SS cilia tips, we found two classes with expected 16 nm repeat for the *doublet withODAs* section. For SS *doublet noODAs*, we reproducibly detected 3 classes (see Supplementary Figure S2), two of which resembled the canonical 16 nm repeat and another with non-canonical, 8 nm repeat. A fourth class contained a small number of noisy particles. Plotting the distribution of classes we found subtomograms with non-canonical IJ more frequently closer to the tip than those with canonical IJ and inspection of the consensus doublet noODAs, non-canonical IJ averages showed it had fewer components bound. We therefore conclude the non-canonical IJ architecture represents an earlier stage of assembly than the canonical IJ. The same workflow was applied to DMTs in Rg cilia tips and the mature axoneme. In Rg cilia tips, classification of the *doublet noODAs* section into two or three classes yielded averages with 8 nm repeat. These appeared similar to the non-canonical IJ architecture obtained in analysis of SS *doublet noODAs*, reinforcing our conclusion that the canonical IJ is generated after the non-canonical IJ. Based on this analysis, we refer to the subtomogram averages of the *singlet* as ‘Stage 1’, *doublet noODAs* with non-canonical IJ as ‘Stage 2’, *doublet noODAs* with canonical IJ as ‘Stage 3’, doublet *withODAs* as ‘Stage 4’ and *mature* as ‘Stage 5’. To compare the densities present across stages, we generated an 8 nm repeat average for Stage 3 (*doublet noODAs*, canonical IJ) by combining both canonical IJ classes. We then performed local, masked refinement of the Stage 2 and 3 averages for all stages. Particle numbers and resolution estimates from RELION PostProcessing for the 8 nm repeat structures are shown in Supplementary Figure S3.

### Generation of DMT 96 nm repeat averages

After generation of 8 nm repeat averages, hierarchical local refinement and classification were performed in RELION 3.1.3 to obtain the 96nm repeat, following a strategy similar to that used in single particle analysis of DMTs ^96^. As illustrated in Supplementary Figure S4, the 8 nm repeat was classified at the IJ to generate a 16 nm repeat average. Subsequent refinement and classification of the seam produced a 48 nm repeat structure. Finally, refinement and classification of the outer A-tubule surface - where large, 96-nm-repeating complexes reside - enabled generation of the full 96 nm repeat structure. Local, masked refinement was performed before each round of 3D classification and a new reference generated from the selected class. T-values were varied between 8 and 24, and the number of classes and size of mask were optimized at each step. Once the 96 nm repeat average was obtained, the particles were re-extracted in a larger box and masked refinement was performed to improve the resolution of key components within the 96 nm repeat (Supplementary Figure S5). Resulting maps were processed using RELION PostProcessing and filtered to their estimated resolution. Composite maps were generated in ChimeraX using volume maximum. This workflow was successful for DMT sections exhibiting canonical 16 nm repeating architecture at the IJ (SS *doublet noODAs* with canonical IJ; SS *doublet withODAs*, Rg *doublet withODAs, mature*). Focused classification of the seam was attempted on the 8 nm repeat averages for SS and Rg Stage 1 and 2, but reliable 48 nm repeat averages could not be obtained. Classification around protofilament A02 at the expected lattice gap site was performed on the 8 nm repeat averages for SS Stage 3 and 4, yielding 96 nm repeat structures containing the lattice gap. The same procedure applied to SS Stage 1 and 2 did not resolve the lattice gap.

### Generating an integrated model of the DMT assembly process

To generate a model of DMT assembly, we integrated data from SS and Rg subtomogram averages. Information from all available periodicities was used; 8 nm averages for Stages 1-4 in SS and Rg conditions and Stage 5 and 96 nm repeat averages for SS Stages 3-4, Rg Stage 4 and Stage 5. The 8 nm repeat averages enabled comparison of A-tubule MIPs, B-tubule MIPs and coiled-coil MAPs (ccMAPs) across all assembly stages (see Supplementary Figure S6). These averages were used to assess the occupancy grooves between protofilaments (visible in cross-sectional projections), and components with 8 nm periodicity (e.g. IJ filament, A-tubule luminal MIPs). The density for some components with longer periodicity (e.g. the seam binding components) were also visible in the 8 nm averages, although their density was subsampled and therefore weaker. Using the 96 nm repeat averages, we assigned the binding of molecular components visible at the resolution obtained (23-33 Å). We fit the PDB 8GLV into our maps after filtering them to the same resolution and adjusting their threshold so microtubule signals were matched. We assigned the components to an assembly stage with two levels of granularity. Per-subunit assignment was given for components tightly associated to the microtubule lattice (MIPs and MAPs). Complex-level assignment was given for larger protein complexes on the DMT exterior, based on the presence of density in the expected position for each complex. For the single-headed IDAs, density was assessed at the microtubule attachment site since the motor domains appeared flexible. For RSPs, density was assessed at the bases rather than the heads, which lie further from the DMT surface. In cases where discrepancies were observed between SS and Rg Stage 3-4 components, we introduced intermediate stages (Stage 3.5 or 4.5) to indicate the components can also be added later in assembly. We interpret these differences as reflecting the same underlying assembly process, with Rg tips exhibiting more spatially extended early-stage features due to active assembly. Partial or weak density in the 96nm repeat averages were also noted and taken into account. A summary of our observations is tabulated in Supplementary Figure S7 and components assigned as present in each stage provided in Supplementary Figure S8.

### Subtomogram averaging of the MT capping complexes

IMOD models were generated with scattered points placed at the center of each MT-end capping structure and these were masked as indicated in Supplementary Figure S9. The models were converted into RELION star files using the same workflow as for DMTs and subtomograms extracted at bin 4 (14.88 Å/pix). The A-tubule plugs were assigned the same index as the DMTs, allowing initial angles to be assigned by transferring the median angles from the *singlet* section on a per-MT basis. Similarly, Ring 1 in the CMC was given prior angles from the LLS section in each tomogram. Ring 3 was flexibly attached to the CP end and so initial tilt and psi angles were manually assigned. In each case, local refinement was performed in RELION 3.1.3. We found Ring 3 had 9-fold symmetry and re-refined this structure with C3 symmetry.

### Subtomogram averaging of the central apparatus

To analyze the assembly of the CA at the ciliary tip, we traced the central path of the CP microtubules from their plus ends. An overview of this process is shown in Supplementary Figure S10. Subtomograms were extracted from bin4 tomograms using the same approach as for DMTs, with sampling every 8 nm along the CP in the four regions defined during initial modelling (LLS, *Bare, Zone 1* and *Zone 2)*. Tilt and psi angles were derived from the microtubule path in the tomograms and used as priors during refinement. However, global refinement of all CP particles did not reliably distinguish between the C1 and C2 microtubules, resulting in misaligned rot angles and poor averages. Therefore, approximate rot angles were manually assigned on a per-CP, per-section basis via visual inspection in IMOD. Initial masked refinement of the entire CA showed the LLS is not tightly associated with a specific site on the CP microtubules. In the *Bare* and *Zone 1* regions, the C1 and C2 protofilaments appeared smeared, prompting separate masked refinement of each microtubule and its associated projection arms. This analysis showed that C1 and C2 maintain a fixed relative position only in *Zone 2*. Inspection of these averages showed *Zone 1* had the C2a arm on the surface of C2 while *Zone 2* had all four major projection arms. We did not find evidence of repetitive proteins binding the *Bare* region. To identify components present in *Zone 1* and *Zone 2*, we recentered the subtomograms on each microtubule. Classification based on the surface densities yielded a 16 nm repeat structure for C2 in both zones and a 32 nm repeat structure of C1 in Zone 2 (see Supplementary Figure S11,S12). Similar as for DMTs, tip averages were locally refined and compared to *mature* CA structures obtained from non-tip tomograms. Due to having ∼9-fold fewer subtomograms than for DMTs, we performed analysis of the CA architecture only in the SS tips.

### Generating a model for CA assembly

To construct a model of CA assembly, we compared the components present in the 16 nm repeat averages of *Zone 1* and the 32 nm repeat averages of *Zone 2* to the *mature* CA structure. Structural modelling was performed using a composite of PDB 7SOM ^1^ and PDB 7N61^2^ for C2 and PDB 7SQC^1^ and PDB 7N6G ^2^ for C1. The motor arm is flexible but was visualized in Zone 2 through classification of density near the C2a arm and masked refinement of the C2b region (see Supplementary Figure S13). Subtle differences were observed at the periphery of the C2a arm in *Zone 1* compared to *Zone 2* and the *mature* structure. These regions correspond to the positions of FAP70 and FAP147. However, since the density for these components at the base of the C2a arm were consistent across all stages, we attribute the observed difference to increased flexibility rather than absence of these components. Details of our observations are provided in Supplementary Figure S14.

### In silico screening of interaction partners of KLP1

To identify potential interaction partners of KLP1, sequences for all components assigned as present in the Zone 1 average were retrieved from UniProt. Protein-protein interaction predictions were performed using AlphaFold2.3 via AlphaPulldown v1.0.4^97^. The Uniprot IDs used were: KLP1 - A8I9T2; PF20 - P93107; FAP70 - A0A2K3DKW3; FAP65 - A0A2K3DLJ2; FAP20 - A8IU92; FAP196 - A0A2K3CQT7; FAP178 - A0A2K3D8Z6; FAP174 - A8I439; FAP147 - A0A2K3DUG8. Interaction predictions were performed in pulldown mode, with KLP1 as the bait protein. Among the candidates, only PF20 showed a predicted interaction, with inter-chain predicted aligned error (PAE) values below 5 Å consistent across the 5 models generated. The next best scoring protein being FAP147, which had inter-chain PAE score below 10 Å. All other candidates had inter-chain PAE values exceeding 50 Å, indicating a lack of interaction. A hairpin helix within KLP1 and the flexible N-terminus of PF20 were predicted to interact (Fig S10) and the predicted model of their interaction had an iPTM score of 0.74 and mpDockQ/pDockQ scores of 0.45. Ribbon plot was generated using AlphaFold2 predictions of individual full-length structures in SSDraw ^98^.

### Measurement of interprotofilament angles

Interprotofilament angles were measured using the consensus 8 nm repeat subtomogram averages for SS and Rg Stages 1-4. Each map was filtered to its resolution estimated in RELION PostProcessing and opened in ChimeraX. Short protofilament segments, consisting of three tubulin dimers, were manually placed around the circumference of the A-tubule by fitting them into the density using the ChimeraX ‘fit’ command. The rotation between adjacent protofilaments was then measured (using ‘measure rotation’), providing the angular displacement between tubulin dimers in neighboring protofilaments. Measurements were made at three different thresholds for each map (0.7, 1.0, 1.4) with the mean plotted.

### 3D molecular visualization of ciliary tip and assembly process

For 3D visualization of the tip architecture as in Figure 1, subtomogram averages were repositioned using the ArtiaX plugin in ChimeraX^99^. Membrane segmentation was performed using MemBrain-Seg ^100^. Structures without averages (e.g. the net) were visualized using meshes exported from IMOD. The 3D molecular animations in Supplementary Movies 1 and 2 are based on molecular structure files used for mapping of the component presence (PBD 8GLV for the DMT, a composite of PDBs 7SOM and 7N61 for C2 and PDBs 7SQC and 7N6G for C1) and were complemented by subtomogram averages from this study when models were missing - respectively the microtubule capping structures, end plugs and 8 nm repeating densities at the IJ for the DMT. Density for C2b-c-d motor arm was modelled using EMD-24191. The molecular models and subtomogram averages were converted to polygonal meshes using ChimeraX and imported into Autodesk Maya for animation, where a few additional structures were modelled using tomograms as reference, respectively the hairs at the tip of the A-tubule plug in the case of the microtubule doublet, and the CMC and LLS in the case of the central pair of microtubules. Key assembly stages were defined based on evidence shown throughout the manuscript and summarized in Supplementary Methods 1 and 2 and animated using keyframe interpolation. The animation was rendered as a series of images and final compositing was done using Adobe After Effects. Still images were rendered to highlight that relevant MIPs can fit into the hole created in the microtubule by the transient removal of one tubulin dimer (shown in Fig S4). The regions missing from the corresponding molecular models are all predicted to be disordered by AlphaFold, and we thus assumed that they would also fit into the hole.

### Orthology inference

A total of 27 reference proteomes were downloaded from UniProt (https://www.uniprot.org/proteomes). The proteomes considered include representatives of all major eukaryotic groups (Protist, Metazoa, Plants, and Green Algae), ciliated and non-ciliated model organisms, and 4 Chlamydomonadales species (Supplementary Table 1). Orthologues of MIPs and MAPs in the *C. reinhardtii* DMT (Supplementary Table 2) were inferred between species using OrthoFinder (v.2.3.7) ^101^. Orthologous groups (orthogroups) were identified across all 27 species using the full reference proteomes. Orthogroups containing *C. reinhardtii* ciliary proteins were identified and used for comparative analyses. All 46 of the *C. reinhardtii* proteins were assigned to an orthogroup, and a total of 841 orthologous proteins (Supplementary Table 3) were identified across the 27 organisms. Individual protein searches against reference proteomes were performed using the function phmmer in HHMER v3.4 (http://hmmer.org/).

### Figure generation

Figures were generated in ChimeraX 1.7.1. Graphs were plotted in GraphPad Prism.

## Supplementary Information

**Figure S1.**
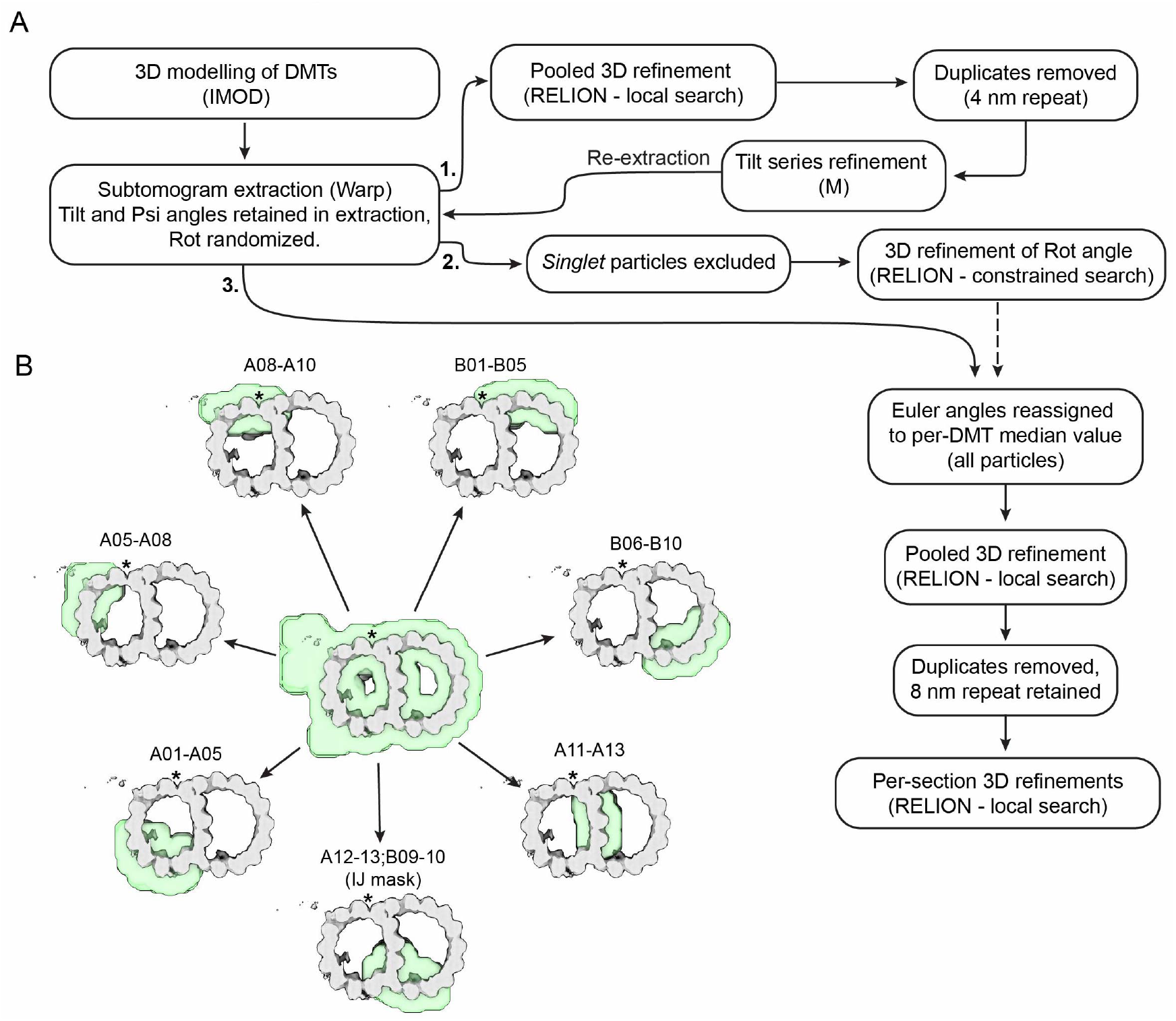
DMT data processing pipeline. A) Workflow used for initial alignment of DMT and generation of 8 nm repeat structures for *singlet,doublet noODAs* and *doublet withODAs* sections from SS and Rg cilia tips. B) Consensus map (grey) and masks (transparent green) used for local refinement of the SS *doublet withODAs* 8 nm repeat averages. Equivalent masks were used for all other sections (*singlet,doublet noODAs*) and conditions (Rg, *mature*)

**Figure S2.**
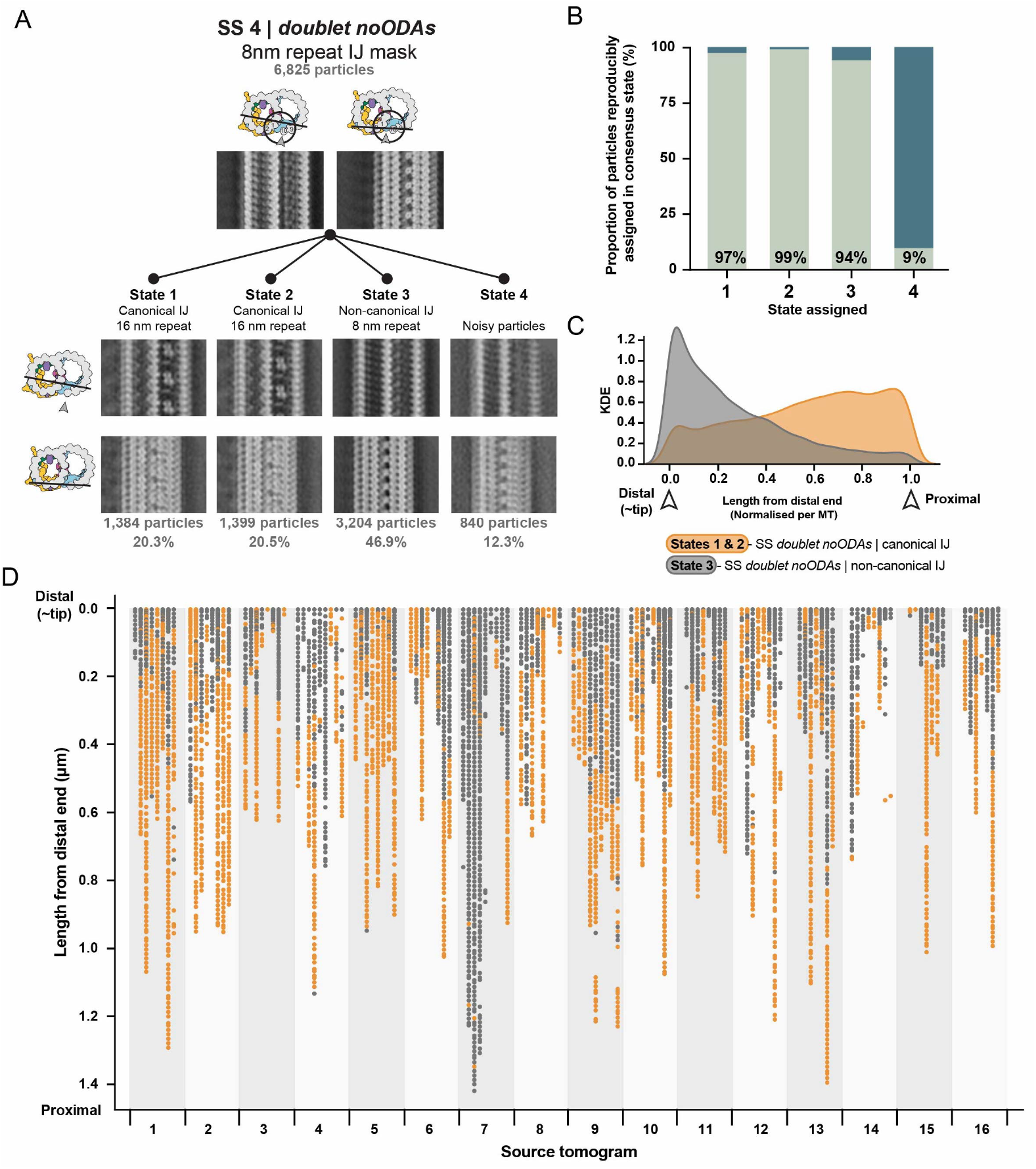
IJ classification in SS *doublet noODAs* and distribution of classes. A) Results of 3D classification of the 8 nm repeat average for SS *doublet noODAs*. Local refinement was performed around the IJ, followed by classification into four classes. Two classes exhibited canonical 16 nm repeat architecture in different registers (States 1 and 2), one class showed a non-canonical 8 nm repeat (State 3), and one class contained noisy or poorly aligned particles (State 4). B) Reproducibility of classification results. The classification procedure was repeated four times. Subtomograms assigned to the same state in at least three out of four runs were considered reproducibly classified. State assignments are as in (A). C) Kernel Density Estimation (KDE) plot showing the spatial distribution of subtomograms with canonical (orange) and non-canonical (grey) IJ architecture. The Euclidean distance of each subtomogram from the tip of its microtubule was min–max normalized per microtubule and data from all tomograms pooled. D) Linear distribution of subtomograms along the length of the DMT. Each point represents a subtomogram, plotted along a line aligned at the microtubule tip. Canonical (orange) and non-canonical (grey) IJ states are shown for SS *doublet noODAs* section. Gaps in the distribution reflect removal of particles assigned to the noisy State 4.

**Figure S3.**
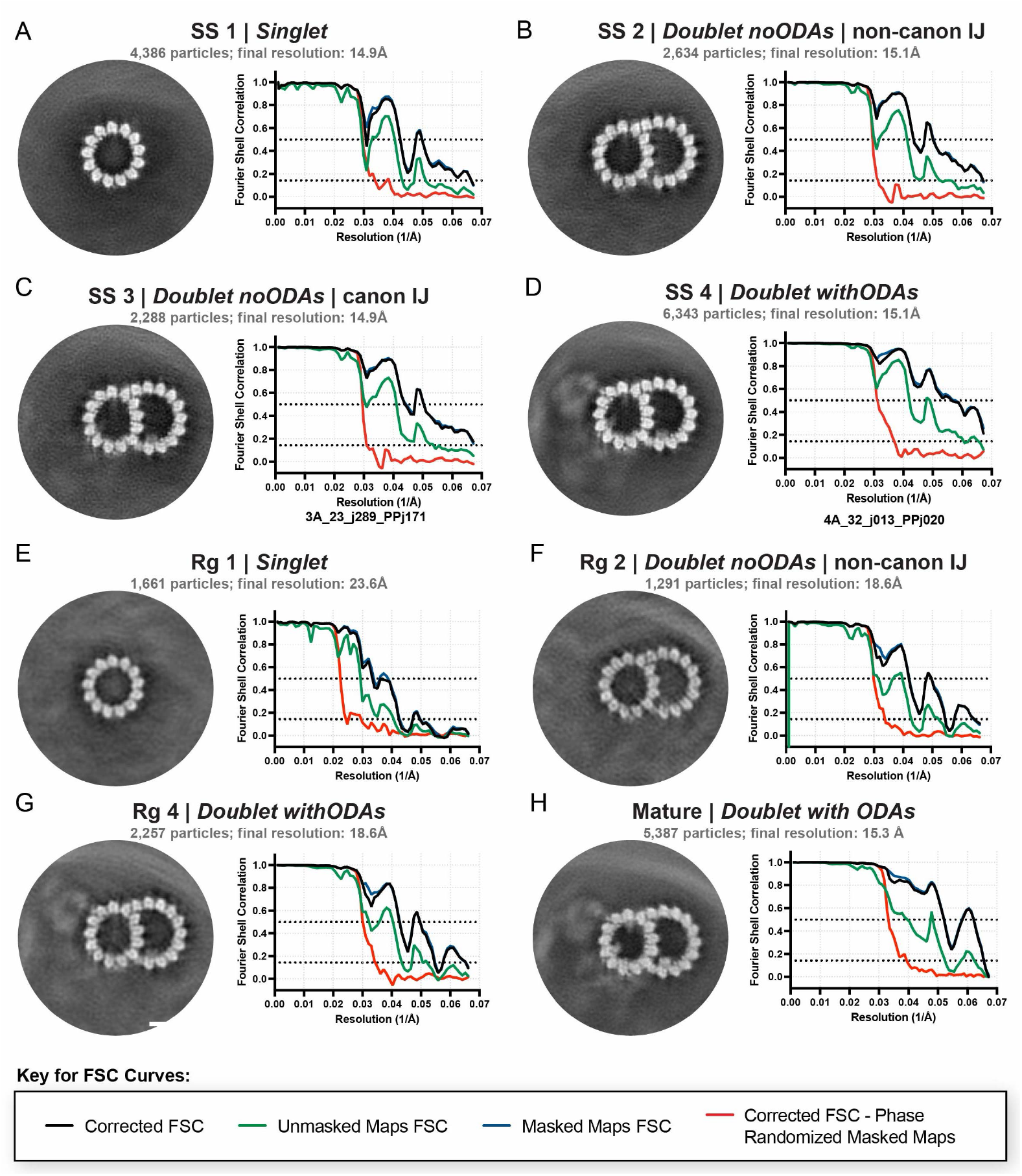
Resolution obtained for consensus DMT 8 nm repeat averages. A-H) For each panel, a projection of the subtomogram average obtained after consensus refinement of DMT sections with 8 nm periodicity is displayed. On the right of each panel, Fourier Shell Correlation (FSC) curves generated in RELION PostProcessing are displayed. Gold-standard resolution estimates are reported at the 0.143 cutoff, along with the number of particles contributing to each average. All refinements were performed using the central mask shown in Methods Supplement 1 - Fig 1B.

**Figure S4.**
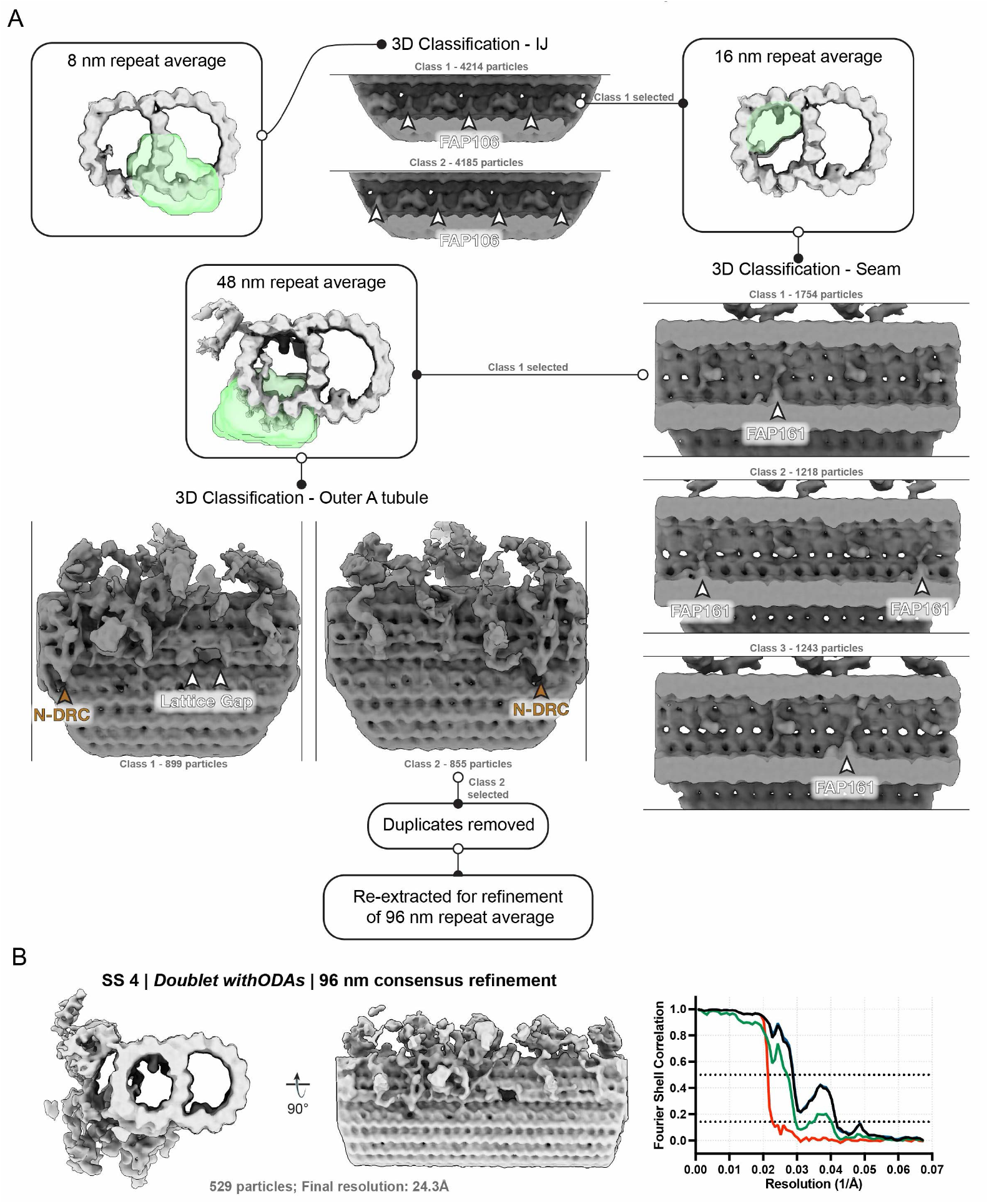
Hierarchical classification of DMTs to obtain 96 nm repeat. A) Hierarchical classification workflow used to generate the 96 nm repeat average for SS *doublet withODAs* sectiion. For each round of classification, the 3D reference (silver), mask (transparent green) and resulting classes (grey) is shown. The same procedure was followed for SS *doublet noODAs*, canonical IJ, Rg *doublet noODAs* and *mature*. B) Consensus map of the 96 nm repeat average for SS *doublet withODAs*. Cross-sectional and longitudinal views are shown along with FSC curve resulting from RELION PostProcessing. The gold-standard resolution estimate at the 0.143 cutoff is indicated, along with the number of particles contributing to the final average.

**Figure S5.**
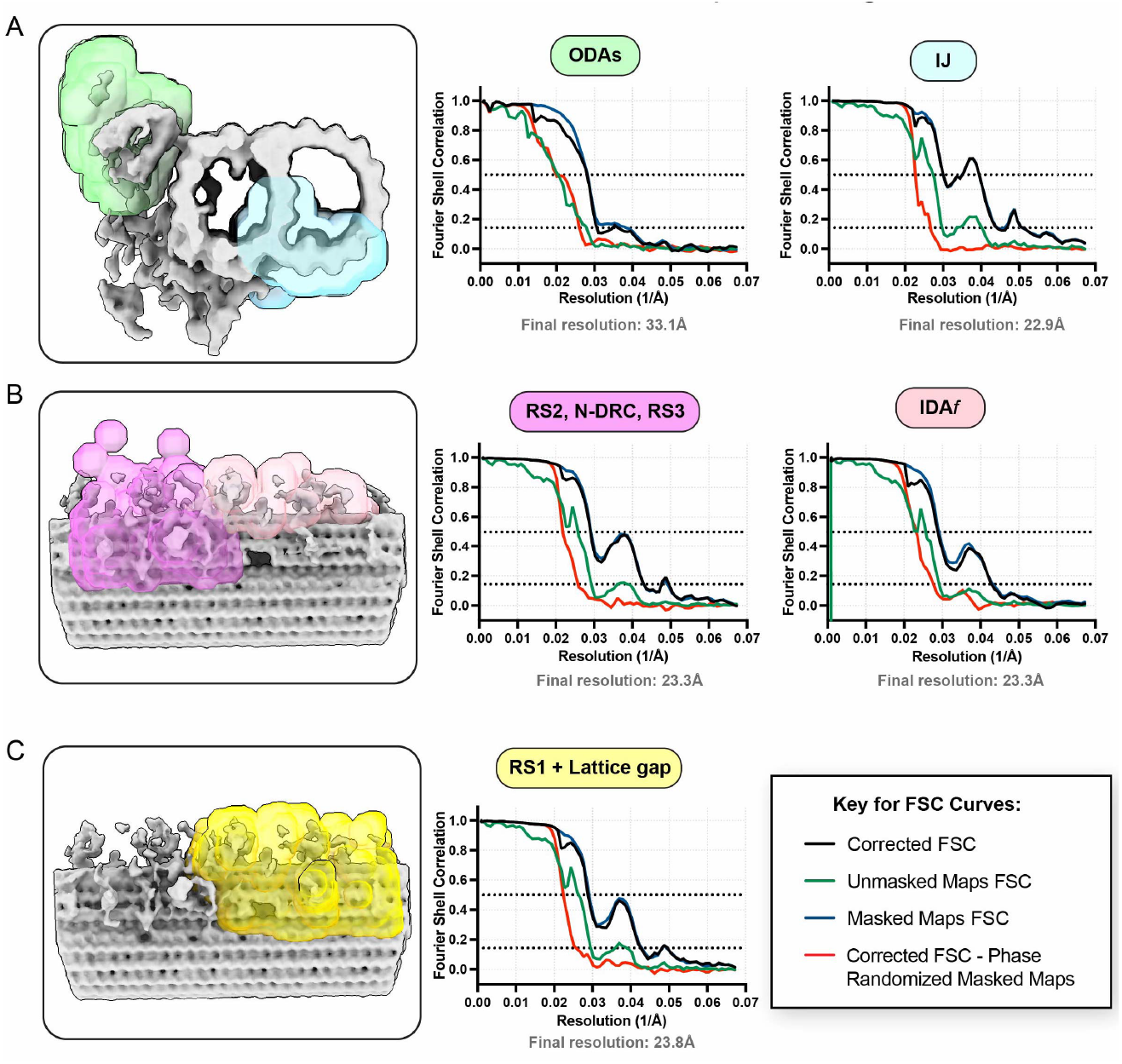
Local refinement of SS *doublet withODAs* 96 nm repeat average. A-C) Consensus map (silver) and masks (transparent, colored) used for local refinement of the SS *doublet withODAs* 96 nm repeat averages. FSC curves for each masked refinement is shown and gold-standard resolution estimates are reported at the 0.143 cutoff. Equivalent masks were used for Rg *doublet withODAs, mature* and SS *doublet noODAs* structures.

**Figure S6.**
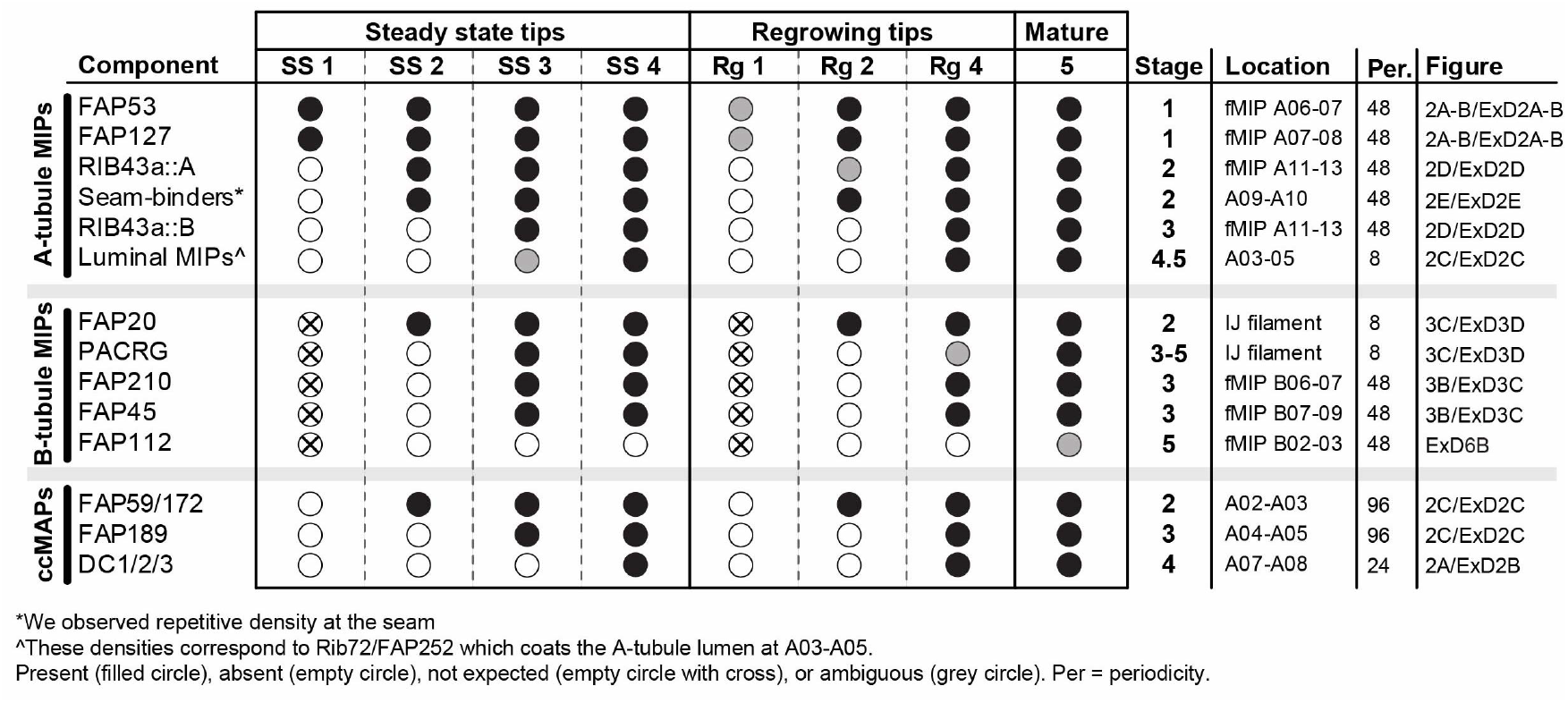
DMT component mapping based on 8 nm repeat averages

**Figure S7.**
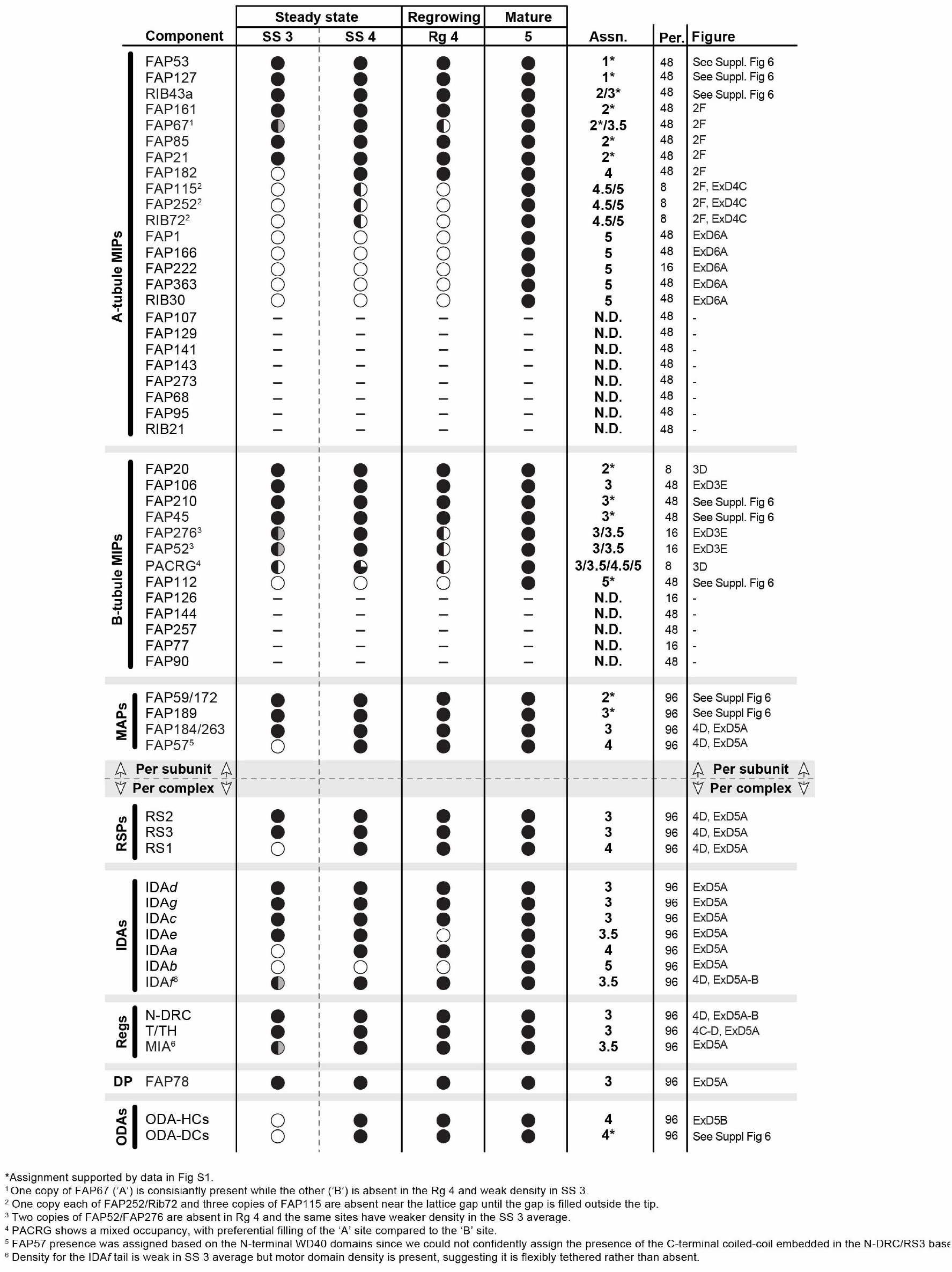
DMT component mapping based on 96 nm repeat averages

**Figure S8.**
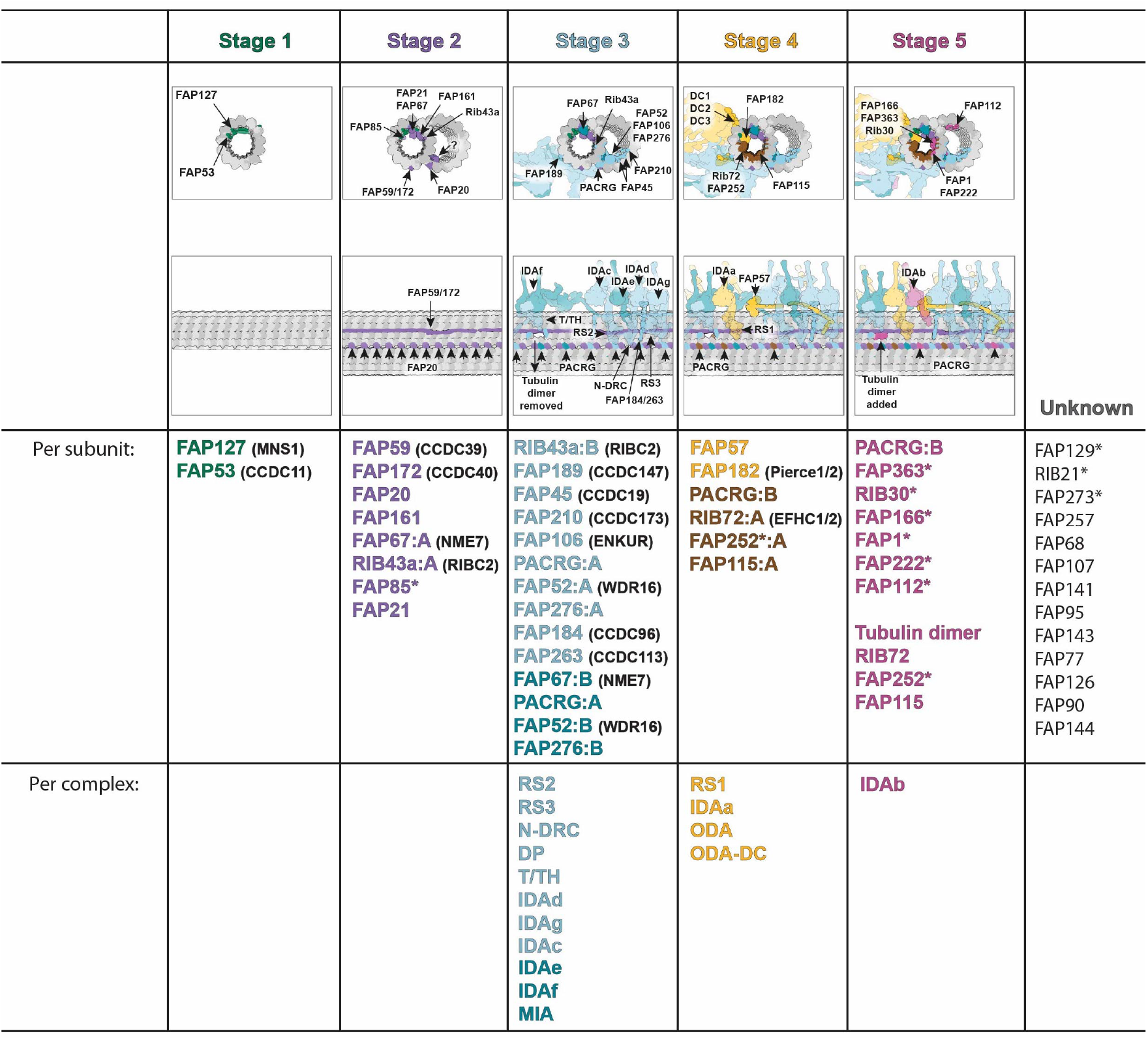
List of axonemal components added at each DMT assembly stage. Proposed model of DMT assembly stages, with components listed by stage of incorporation. Axonemal components were grouped by granularity of assignment; MIPs and MAPs which are tightly microtubule-associated were analyzed per-subunit; external complexes were assessed as complete units. Names of mammalian homologues are given in brackets where available. Asterisks indicates proteins which lack structural homologues outside of Chlamydomonas (based on PMID:37327785).

**Figure S9.**
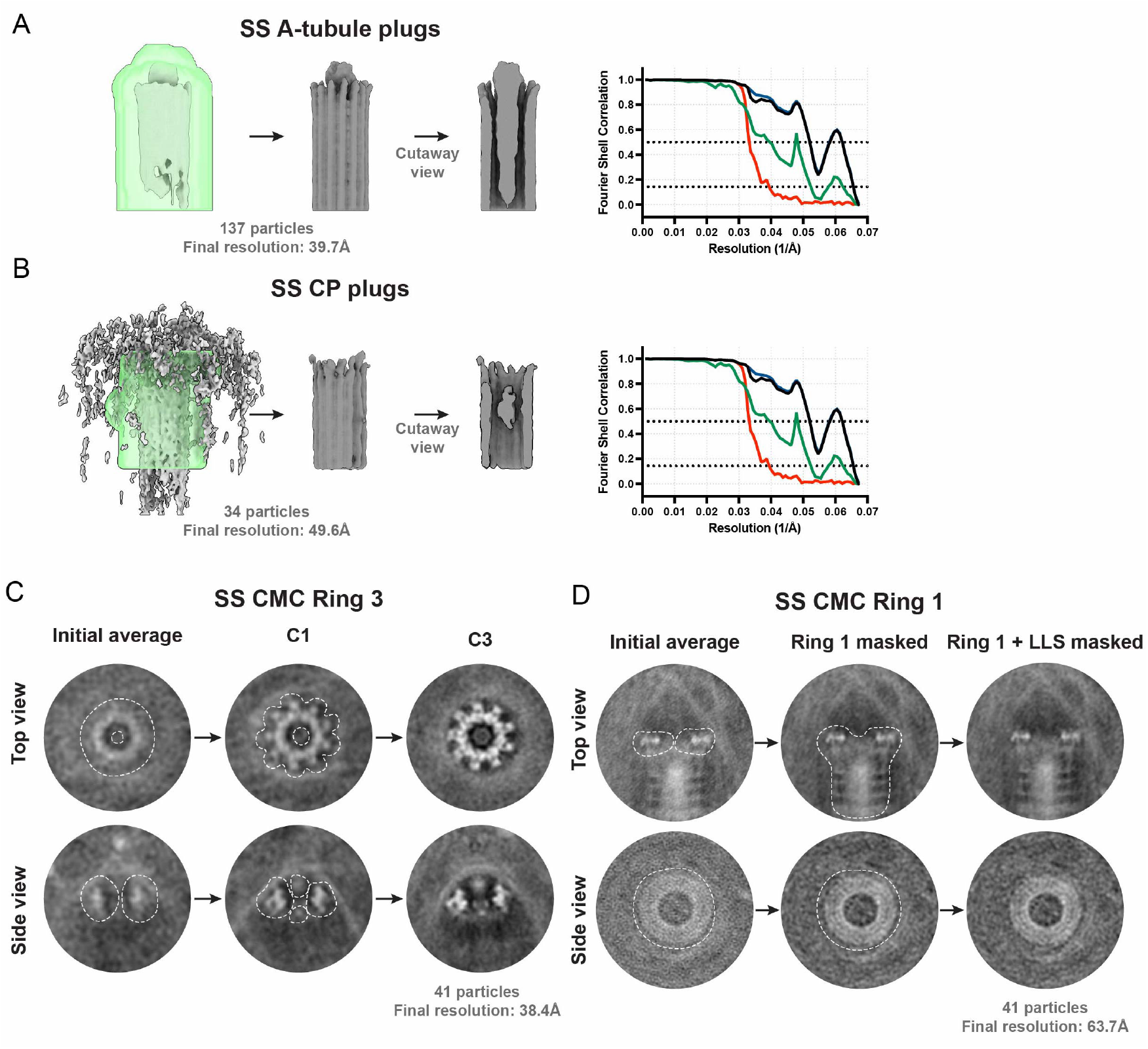
Subtomogram averaging of MT end capping complexes. A) Subtomogram average of A-tubule plugs from SS cilia. Initial average is far left, with mask in transparent green and resulting average in center. Cutaway view shows extent of plug-like structure in microtubule lumen. FSC curve resulting from RELION PostProcessing is shown. B) As in A), for CP plugs. C) Projections of subtomogram average of membrane-adjacent Ring 3 in the CMC of SS cilia tips. Top and side views are shown, with initial average on left, locally refined average with C1 symmetry on right ad locally refined average with C3 symmetry shown on right. Mask position is indicated with dashed line. D) Projections of subtomogram average of microtubule-end-associated Ring 1 in the CMC of SS cilia tips. Initial average with manually assigned angles is shown on the left, local refinement where only Ring 1 is masked is in center and refinement where Ring 1 and the visible portion of the LLS is present is on the right.

**Figure S10.**
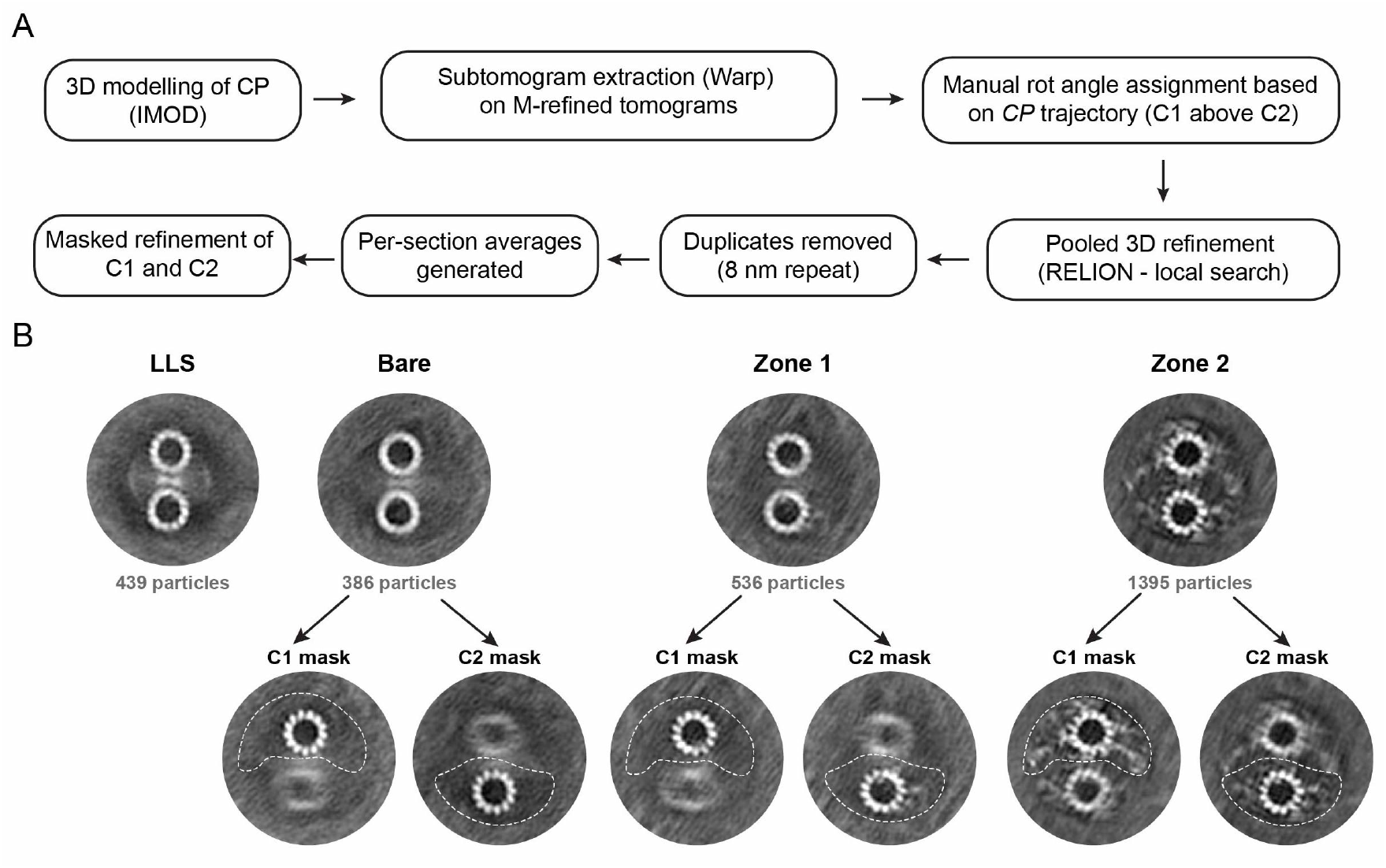
CA data processing pipeline for 8 nm repeat average structures. A) Workflow used for initial alignment of CP and generation of 8 nm repeat structures for LLS, Bare and *Zone 1* and *Zone 2* sections from SS cilia tips. B) Projections of subtomogram averages for each CP section. Top panels show results from local refinement when entire CA is masked, lower panels show averages resulting from when either the C1 or C2 was masked separately. Mask positions are indicated with dotted lines.

**Figure S11.**
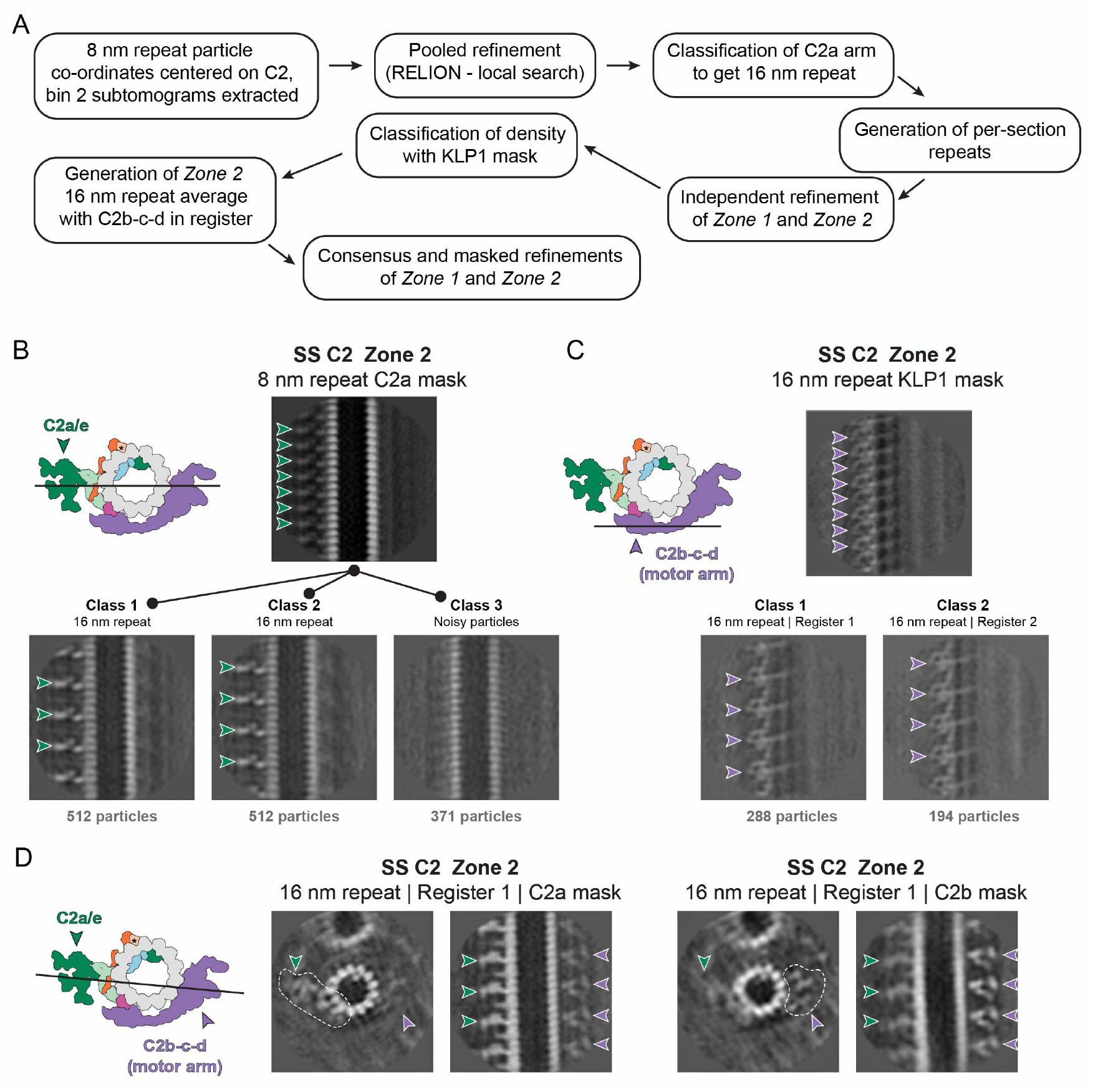
Classification workflow for C2 microtubules. A) Workflow used for generation of 16 nm repeat structures for *Zone 1* and *Zone 2* in SS cilia tips. This resulted in obtaining the 16 nm repeat average for *Zone 1* and the 16nm repeat average for *Zone 2* with the C2b-c-d motor arm in register. The same workflow was used for subtomograms from the mature axoneme, except the pooled average was not generated prior to classification. B) Classification of the C2a arm to generate the 16 nm repeat average for *Zone 2* in SS cilia tips. The C2a arm was masked and refined prior to classification. Viewing position is indicated by cartoon. C) Classification of the C2b-c-d motor arm close to the KLP1 array to generate the 16 nm repeat average of *Zone 2* with C2b-c-d in register. The same procedure was performed for the *Zone 1* average but the classes were indistinguishable and contained noise in the target region. D) Subtomogram averages of Class 1 generated in C) showing the result when the C2a arm (left) or C2b-c-d motor arm is masked, showing both are present in Zone 2.

**Figure S12.**
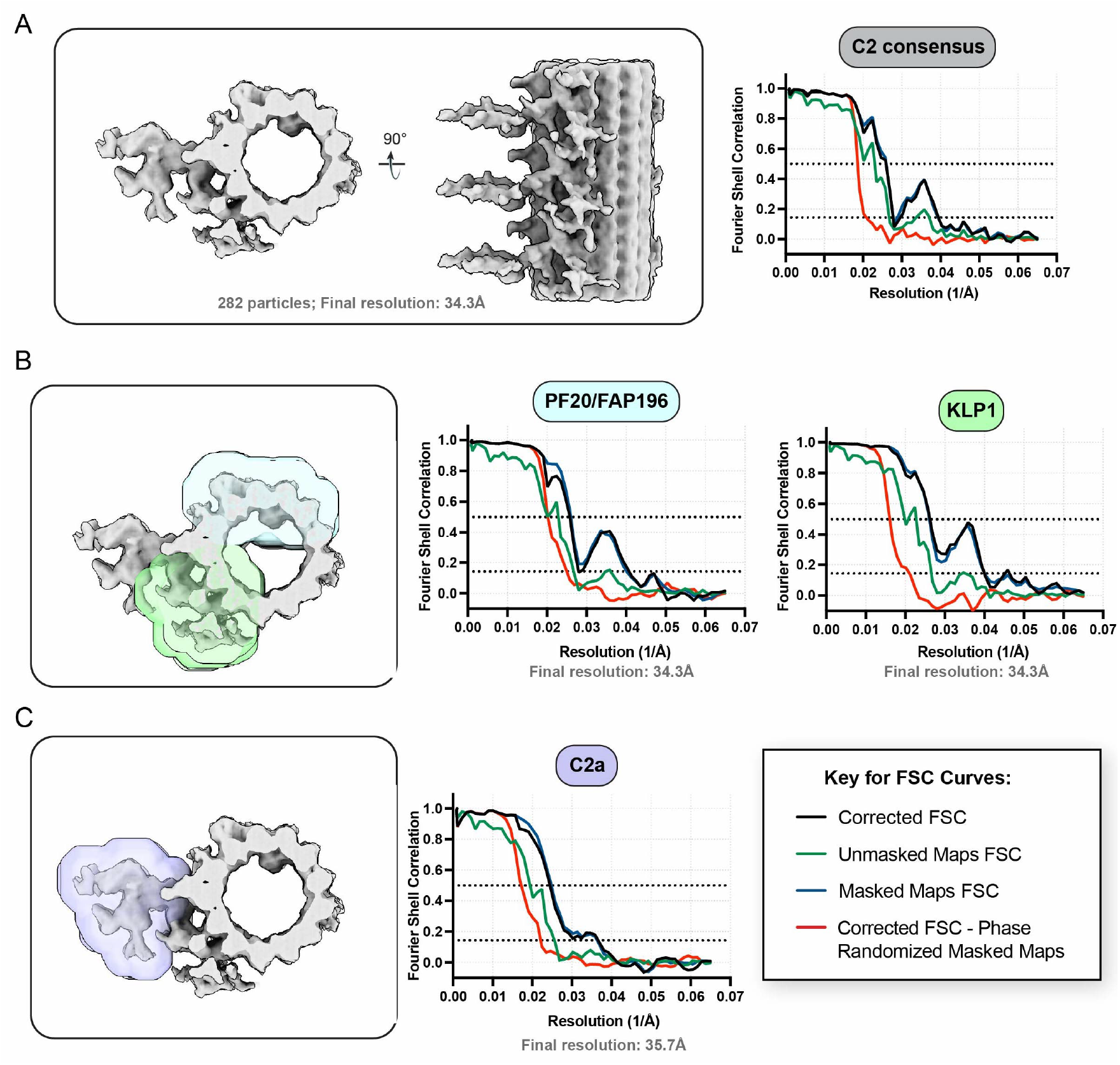
Local refinement of C2 microtubules. A-C) Consensus map (silver) and masks (transparent, colored) used for masked refinement of the SS *Zone 2* 16 nm repeat averages with KLP1 register 1. FSC curves for each masked refinement is shown and gold-standard resolution estimates are reported at the 0.143 cutoff. Equivalent masks were used for *Zone 1* and *mature* C2 16nm repeat structures.

**Figure S13.**
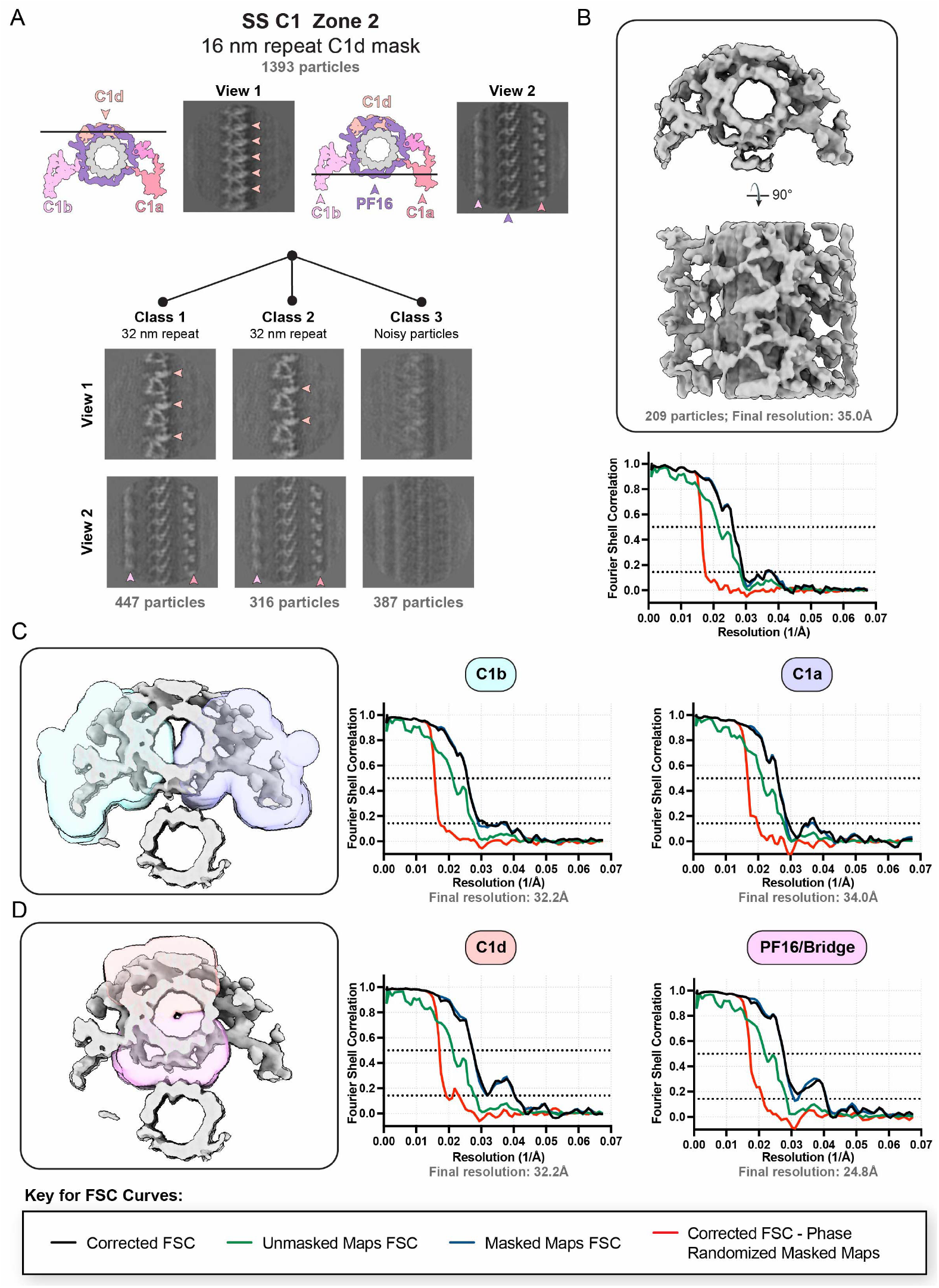
Classification to generate 32 nm repeat structure of the C1 microtubule. A) Projections of subtomogram averages showing classification used to obtain 32 nm repeat average of C1 in SS Zone 2. C1d was masked for refinement and classification. Class 1 was selected and duplicates removed before consensus refinement. B) Surface view of consensus refinement for C1 in SS *Zone 2* 32 nm repeat average. Top and side views are shown, along with FSC curve. gGld-standard resolution estimates are reported at the 0.143 cutoff. C-D) Consensus map (silver) and masks (transparent, colored) used for local masked refinement of the SS *Zone 2* 32 nm repeat averages.

**Figure S14.**
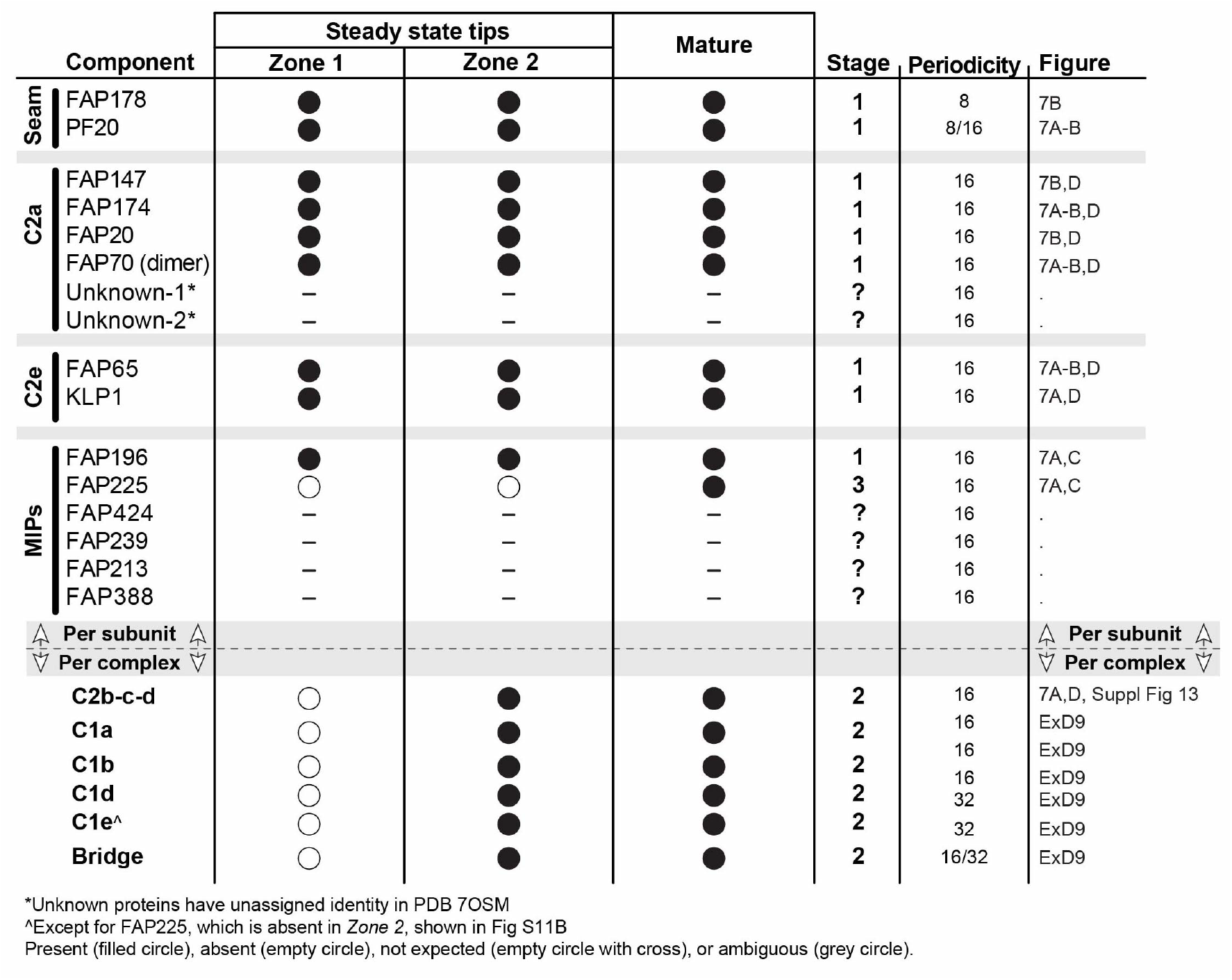
Data used for generating CA assembly model

